# 12/15-Lipoxygenase orchestrates murine wound healing via PPARγ-activating oxylipins acting holistically to dampen inflammation

**DOI:** 10.1101/2025.03.26.645456

**Authors:** Christopher P Thomas, Victoria J Tyrrell, James J Burston, Sam R C Johnson, Maceler Aldrovandi, Jorge Alvarez-Jarreta, Rossa Inglis, Adam Leonard, Lydia Fice, Jeremie Costales, Stefania Carobbio, Antonio Vidal-Puig, Majd Protty, Carol Guy, Robert Andrews, Barbara Szomolay, Ben C Cossins, Ana Cardus Figueras, Simon A Jones, Valerie B O’Donnell

## Abstract

12/15-lipoxygenase (12/15-LOX, *Alox15*) generates bioactive oxygenated lipids during inflammation, however its homeostatic role(s) in normal healing are unclear. Here, the role of 12/15-LOX in resolving skin wounds was elucidated, focusing on how its lipids act together in physiologically relevant amounts. In mice, wounding caused acute appearance of 12/15-LOX-expressing macrophages and stem cells, coupled to early generation of ∼12 monohydroxy-oxylipins and enzymatically oxygenated phospholipids (eoxPL). *Alox15* deletion increased α-smooth muscle actin, collagen deposition, stem cell/fibroblast proliferation, IL6/pSTAT3, pSMAD3, and IFN-γ levels. Conversely, CD206 expression, F480+ cells, MMP9 and MMP2 activities were reduced. *Alox15^-/-^* skin was deficient in PPARγ/adiponectin activity. Furthermore, while pro-inflammatory genes were upregulated as normal during wounding, many including *Il6, Il1b, ccl4, Cd14, Cd274, Clec4d, Clec4e, Csf3,* and *Cxcl2* failed to revert to baseline during healing, indicating disruption of an anti-inflammatory brake. Reconstituting *Alox15^-/-^* wounds with a physiological mixture of *Alox15*-derived primary oxylipins generated by healing wounds restored MMP and dampened collagen deposition. The oxylipin mixture activated PPARγ *in vitro*, while *in vivo,* the PPARγ co-activator, *Helz2*, was significantly upregulated. Additional inflammatory and proliferative gene networks impacted by *Alox15^-/-^*included *Elf4, Cebpb* and *Tcf3*, with many of their associated genes significantly dysregulated. In summary, the impact of 12/15-LOX is ascribed to the deficiency of abundantly generated monohydroxy oxylipins acting together via PPARγ/adiponectin. The identification of multiple gene alterations reveals several new targets for treatment of non-healing wounds. Our studies demonstrate that abundant 12/15-LOX oxylipins act together, dampening inflammation *in vivo*, revealing a need to consider lipid signaling holistically.

**Significance statement:** Defective wound healing is a significant global clinical problem. Macrophage 12/15-lipoxygenase (12/15-LOX, *Alox15*) generates abundant lipid mediators termed oxylipins during inflammation. However, its physiological role during resolving wound healing is unclear, with studies so far assessing the bioactivity of individual lipids pharmacologically, rather than holistically in physiological amounts. Here, we report that *Alox15* deficiency in mice caused a fibrotic response with failure to dampen inflammation, due to a dysregulated PPARγ/adiponectin axis. Treatment of *Alox15^-/-^* wounds with physiological mixtures of PPARγ-activating 12/15-LOX primary monohydroxy products restored the phenotype. Several transcriptional networks (*Elf4, Cebpb* and *Tcf3*) controlled by *Alox15* were uncovered, identifying new targets for promoting physiological wound healing.

## Introduction

12/15-LOX (*Alox15*) is a leukocyte enzyme highly expressed in murine resident peritoneal macrophages. The human homolog, 15-LOX1 (*ALOX15*) is inducible in peripheral monocytes in response to Th2 cytokines, and expressed basally in reticulocytes, eosinophils and airway epithelium(1, 2). *Alox15^-/-^*mice are protected against atherosclerosis, diabetes, hypertension and abdominal aortic aneurysm, and show reduced thrombosis, while conversely, they develop worse arthritis(3–7). This indicates that the pathway is a significant player in inflammatory vascular disease. However, while central roles in disease are established, the function of 12/15-LOX in normal healing is less clear.

12/15-LOX generates families of structurally related lipid mediators through oxidation of unsaturated fatty acids (FA) and complex lipids, including phospholipids (PL) and cholesteryl esters (CEs)(8–11). The monohydroxy forms of oxidized FAs are first generated by LOXs, with the most abundant being usually derived from arachidonate (AA). These 12/15-LOX derived lipids can independently mediate bioactions relevant to inflammation, such as activation of PPARγ (which dampens cytokines such as IL6 and TNFα)(7, 12–20). Studies up to now generally focused on their bioactions when added individually, for example(21–24). However, *in vivo* they are not generated in isolation but in mixtures comprising large numbers of species at varying amounts. This is particularly relevant to PPARγ, which recognizes overall ligand “tone” at relatively low affinity, rather than specific lipid structures at high affinity via GPCRs. The most quantitively abundant free acid 12/15-LOX products are monohydroxy FAs, from arachidonic acid (AA) and other polyunsaturated fatty acids. Additionally, “specialized pro-resolving mediators” (SPM), such as resolvins, protectins, and maresins are described as rarer products of the pathway(25). Here, the primary monohydroxy FAs are further metabolized, generating oxygenated di- and tri-hydroxy FAs reported to signal via activation of GPCRs that include ALX/FPR2, DRV1/GPR32, DRV2/GPR18, and ERV1/ChemR23(26, 27). However, while SPM can dampen inflammation pharmacologically, their endogenous generation and GPCR binding were recently queried(28–34).

Skin wounding (punch biopsy) represents a tractable model of physiological inflammation resolution that includes four phases: hemostasis, inflammation, proliferation, and remodeling, representing an ideal model in which to test the impact of *Alox15*. Throughout this, lymphoid, myeloid, and tissue-resident cells interact, producing signaling molecules which work in an orchestrated manner. During hemostasis, clotting factors and angiogenic factors decrease bleeding and stimulate formation of new blood vessels(35). During the inflammatory phase, neutrophil and macrophage infiltration supports release of chemokines and cytokines, inflammatory agents and antigen control factors(36). Later, the proliferation phase is characterized by fibroblast and keratinocyte migration from the wound edge, mediating contraction and closure(37). Last, during remodeling, increased deposition and cross-linking of collagen takes place, balanced with removal of excess extracellular matrix by myofibroblast-derived collagenases called matrix metalloproteinases (MMPs)(38). Herein, we used genetic, transcriptomic and lipidomic approaches to determine the role of *Alox15* and its lipids in physiological skin wound healing. We found that the gene plays a critical role in ensuring that the healing response is finely tuned, to enable effective healing. Without 12/15-LOX, cellular and tissue responses proceed at accelerated rates suggestive of fibrosis. Abundant monohydroxy FAs, many already known PPARγ ligands, appear responsible for the phenotype when applied in physiological amounts. Our study highlights a central role for *Alox15* in normal healing, defines several new potential targets for promoting healing, and shows the need to consider lipid biology in a holistic manner when delineating cellular signaling roles that drive health and disease.

## Methods

### Animal model

Mice (8-12 weeks old C57/B6/J) were purchased from Charles River UK (Margate, UK), while *Alox15^-/-^* mice were bred in house (F11, C57BL/6J) in isolators. All animal experiments were performed in accordance with the United Kingdom Home Office Animals (Scientific Procedures) Act of 1986, under License (PPL 30/3334). Generating of healing wounds is described in Supplementary Methods.

### Generation of histological tissue sections and staining protocols

At various time points up to 14 days, wounds were harvested, processed and stained either using DAB or fluorescence immunohistochemistry, as described in Supplementary Methods. Collagen was stained using Masson Trichrome, and images acquired and analyzed using microscopy as described in Supplementary Methods.

### RNASeq

Wound tissue dissected from 2 mice (8 wounds in total, 4 wounds per mouse) to generate each sample (n=4/condition) were snap-frozen in liquid N_2_ before being stored at -80 ^0^C. RNA was isolated using RNeasy MinElute Cleanup Kit (Catalogue number 74204 Qiagen, MD, USA), as described in Supplementary Methods. Total RNA was depleted of ribosomal RNA and sequencing libraries prepared with the Illumina®TruSeq Stranded Total RNA Library Prep Gold (Illumina, Inc) kit using TruSeq CD Index Adapters1 (Illumina, Inc). RNA was sequenced using a 75-base paired-end (2x75bp PE) dual index read format on the HiSeq4000 (Illumina, Inc) according to the manufacturer’s instructions, as described in Supplementary Methods.

### Lipid extraction

Wounds were harvested, homogenized as outlined in Supplementary Methods. Internal standards (5 ng each of PC 14:0_14:0 and PE 14:0_14:0) and 5 ul of eicosanoid internal standards were added. Samples were extracted using a solvent extraction (eoxPL) and solid phase extraction (oxylipins) as outlined in Supplementary Methods. Lipids were reconstituted using methanol and stored at -80 °C until LC/MS/MS.

### LC/MS/MS analysis of oxylipins and eoxPL

Lipids were quantified using reverse phase LC/MS/MS as described in Supplementary Methods. Assay parameters are provided in Supplementary Table 3 and(39) for oxylipins and Supplementary Table 5 for eoxPL. For chiral analysis, lipids were separated using a Chiralpak IA-U column (50×3.0 mm, Diacel) in reverse phase mode, with assay parameters as for oxylipins.

### Gel zymography for MMP activity

Wounds were snap frozen then homogenized and analyzed using Novex™ 10% Zymogram Plus (Gelatin)) gels (Thermo Fisher), as described in Supplementary Methods.

### Cell transfection and reporter assays

HEK293 cells were transfected with mouse PPARγ and the *Firefly* luciferase under the control of 3x Ppar Responsive Element (PPRE)(40), as described in Supplementary Methods.

## Results

### Tissue 12/15-LOX is acutely induced by wounding and associated with higher macrophage numbers

The typical architecture of a wild-type mouse punch wound is shown, showing the dermis, wound bed, scab and wound edge (Figure 1 A). Wounding caused a significant increase in 12/15-LOX^+ve^ cells in the skin at 24 hrs (Figure 1 B-D). Most expression was associated with tissue localized F480^+ve^ macrophages (Figure 1 C). 12/15-LOX was also induced in stem cells located at the base of hair follicles adjacent to the wound, but not distal from it (Figure 1 D). This expression pattern suggests that a soluble mediator signaling in response to wounding maybe responsible for induction and that the enzyme is upregulated early post-wounding in both cell types. The total number of F480^+ve^ monocytes/macrophages in the wound at Day 1 was not impacted by *Alox15* deletion, although there was some reduction later, on Days 4 and 7 (Supplementary Figure 1 A,B). In contrast, neutrophil numbers in the sub-endothelial compartment were unaffected by *Alox15* deletion at any timepoint (Supplementary Figure 1 C).

**Figure 1.**
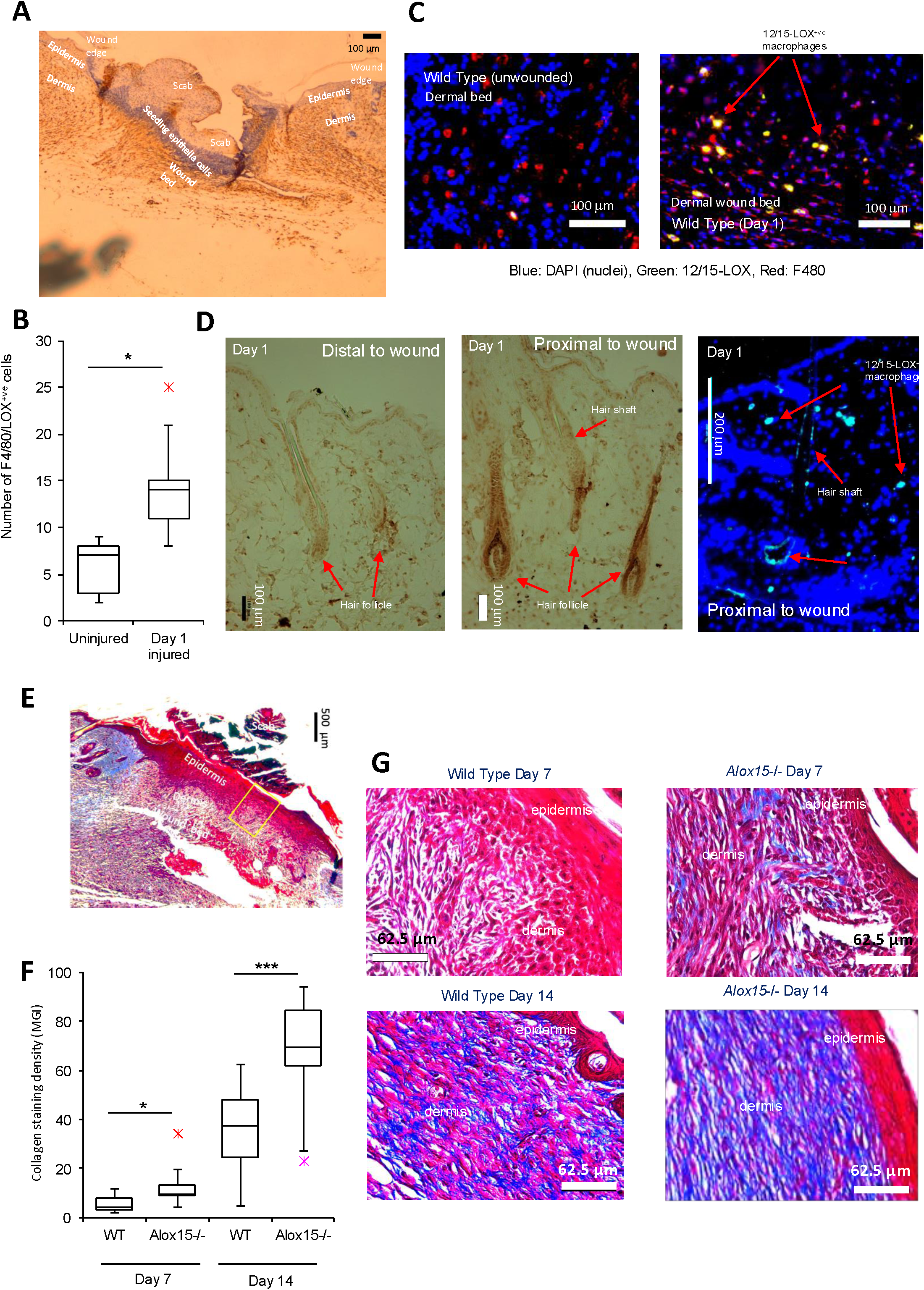
Wounding increases macrophage and hair follicle 12/15-LOX expression, and collagen levels. *Panel A. Representative image of wound architecture. Panel B. Induction of macrophage 12/15-LOX by wounding* 12/15-LOX^+ve^(green)/F4-80^+ve^(red)/DAPI^+ve^(blue) cells were measured at day 1 post wounding (n= 5/group), data were analyzed using one way ANOVA with Tukey post-hoc test * p<0.05, ** p<0.01. *Panel C. Representative images from Panel B*. *Panels D. Expression of 12/15-LOX in hair follicles near the wound edge post-wounding.* Hair shafts are shown on day 1, wild-type mice. The left and center panels show 12/15-LOX DAB^+ve^ staining, and the right panel shows 12/15-LOX^+ve^ green fluorescent staining (with DAPI counterstain). *Panel E. Representative image of wound at low magnification showing region stained for collagen. Panel F. Collagen is elevated in Alox15^-/-^ on days 7 and 14.* Wounds were harvested and analyzed for collagen using Masson’s Trichrome staining (collagen: blue, epithelial cells: deep red, nuclei: black, non-collagen structures: pink) and pixels counted (n = 10-14/group). Unpaired Students t-test, comparing WT and Alox15^-/-^ separately * p<0.05, *** p, 0.005. *Panel G. Representative images from Panel G*.

### Alox15 deletion alters the phenotype of the healing wound, promoting fibroblast stem cell proliferation and differentiation

Healing is characterized by stem cell and fibroblast proliferation, collagen deposition, reconstituting the underlying tissue, as well as epithelial migration and differentiation to form a new covering. Here, smooth muscle actin (myofibroblast marker) and collagen deposition were elevated on *Alox15* deletion (Supplementary Figure 1 D,E, Figure 1 E-G). This was mainly noted during the remodeling phase (day 14), where the majority of the dermal layer in *Alox15^-/-^* wounds was collagen dense (Figure 1 G). Next, we profiled the stem cell marker SSEA3 and the nuclear protein Ki-67, a marker for proliferating cells, in wound beds at day 4. Both were increased in *Alox15^-/-^* with SSEA3 being significantly higher, suggesting that the healing wound at this early stage has a higher number of actively proliferating stem cells (Supplementary Figure 1 F-H). We next determined re-epithelialization of the wound during the inflammatory (day 4) stage using cytokeratins 10 (C10, pink) and 14 (C14, green), which determine epithelial (keratinocyte) cell migration from the wound edge into the bed. Basal keratinocytes which are mitotically active express C14, but during differentiation, they lose C14 and upregulate C10(41). At day 4, the migratory distance of C14^+ve^ and C10^+ve^ epithelial cells into the wound edge was similar for both strains (Supplementary Figure 2 A,B). Overall, the profiles suggest that wound bed keratinocyte differentiation isn’t impacted by *Alox15^-/-^*. Furthermore, the phenotype of non-wounded skin was similar, where in both wild-type and *Alox15^-/-^* mice, C14 is seen to be expressed lower in the epithelium and associated with hair bundle cells, with C10 mainly in terminally differentiated (dead) keratinocytes (corneocytes) on the surface (Supplementary Figure 2 C). Overall, the data indicate that while fibroblast and stem cell differentiation and proliferation in the dermis is impacted, keratinocyte differentiation on the surface of the wound isn’t significantly affected by the absence of 12/15-LOX.

### Elevated TGFβ/IFNγ/IL6 activity is seen in the absence of Alox15

Next, a series of inflammatory pathways were profiled using immunohistochemistry. Protein expression of IL6 was slightly but not significantly higher (Supplementary Figure 2 D), but there was significantly elevated pSTAT3 and pSMAD3 (activated by TGFβ) detected in *Alox15^-/-^* wounds (Figure 2 A,B). Increased IFNγ was also detected, primarily on epithelial cells, while conversely CD206/mannose receptor (a marker of M2 cells), was somewhat reduced (Figure 2 C,D, day 4). Taken together with the collagen, fibroblast and proliferation data, a pro-inflammatory/pro-fibrotic phenotype is suggested for *Alox15* deficiency.

**Figure 2.**
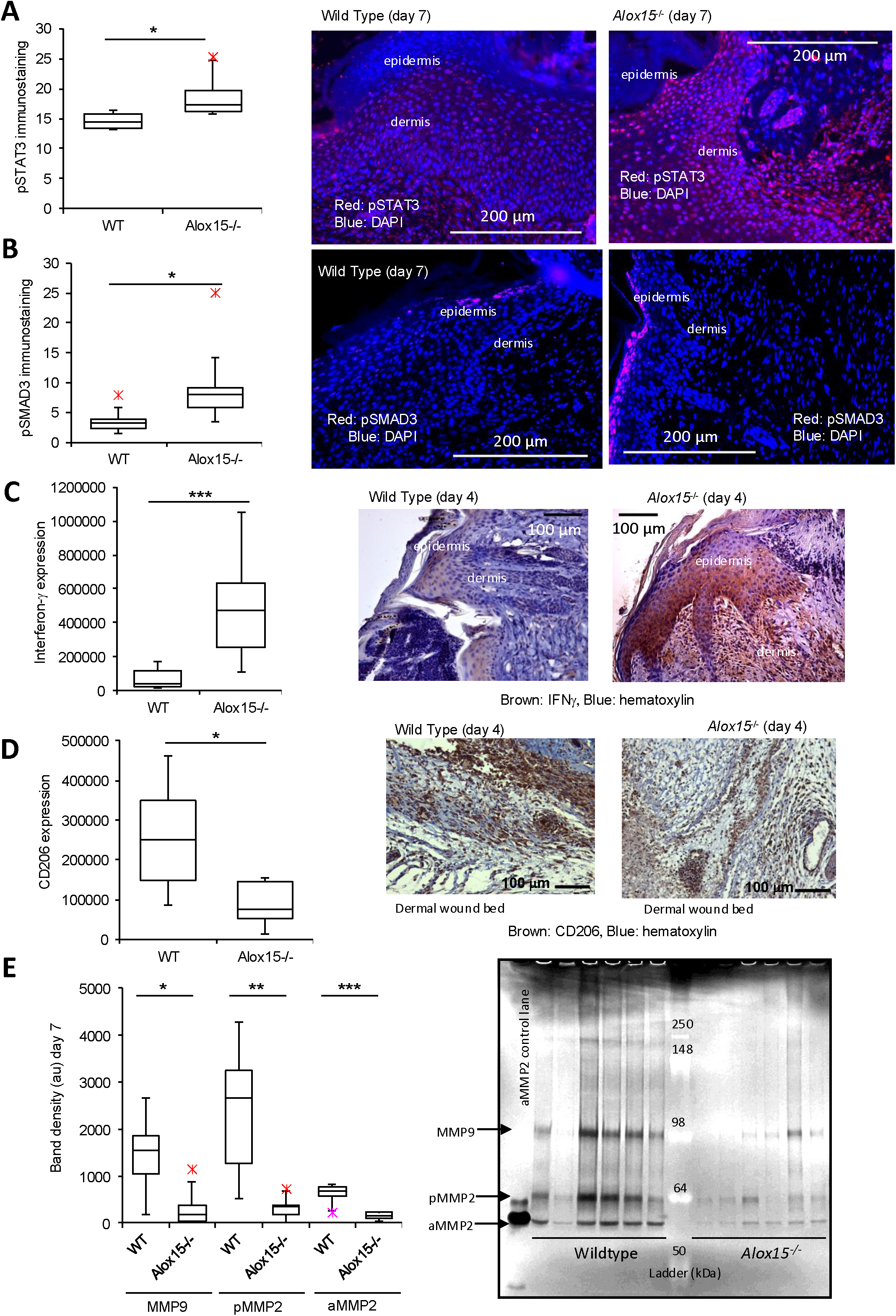
*Alox15^-/-^* wounds show elevated pSTAT3, pSMAD3, and INFγ but decreased CD206 and MMP activities. *Panel A. Alox15^-/-^ wounds show elevated pSTAT3*. pSTAT3 was measured using fluorescence immunohistochemistry. n = 5 - 6/group. *Panel B. Alox15*^-/-^ *wounds show elevated pSMAD3*. pSMAD3 was measured using fluorescence immunohistochemistry. n = 10-11/group. *Panel C. Alox15^-/-^ wounds show elevated IFN*γ. IFNγ was measured using DAB immunohistochemistry. n = 9/group (4-6 fields per wound). *Panel D. Alox15^-/-^ wounds show reduced CD206 expression*. CD206 was measured using DAB immunohistochemistry. n = 5/group (3-6 fields per wound). For all panels data was analyzed an unpaired t-test, * p < 0.05, ** p<0.01. Right panels show representative images for all the proteins analyzed. *Panel E. MMP activities are reduced in Alox15^-/-^ wounds.* MMP activities were measured using zymography. n= 6/group. The ladder shows proteins corresponding to 250, 148, 98, 64 and 50 kDa. Image J was used to calculate the density of each band. The gel is shown (right panel). The impact of *Alox12^-/-^* was analyzed using an unpaired t-test, mean ± SEM, * p < 0.05, ** p<0.01, *** p < 0.005.

### Alox15^-/-^ wounds show reduced matrix metalloprotease (MMP) activities during wound healing

MMPs are collagenases that play a crucial role in regulating the extracellular matrix architecture during wound healing by removing and recycling collagen. Since *Alox15^-/-^* wounds showed elevated collagen deposition, zymography was used to evaluate the activity of critical isoforms, MMP2 (active and pro-forms) and MMP9 on day 7 post-wounding. For all three, collagenase activity was significantly reduced in *Alox15^-/-^*wounds (Figure 2 E).

### The temporal profile of oxylipins is altered by Alox15^-/-^ with many lipids reduced/absent

Using reverse phase LC/MS/MS, ∼100 oxidized fatty acids were profiled in wound tissue, including well-known prostaglandins, thromboxane, eicosanoids, docosanoids and several SPMs. A summary of the 68 lipids detected is shown in a heatmap (Supplementary Figure 3 A). Several monohydroxy lipids were strongly elevated at day 1 but absent in *Alox15^-/-^* wounds (15-HEPE, 14-HDOHE, 17-HDOHE, 13-HOTrE) (Figure 3 A). 12-HETE/12-HEPE, and 15-HETE/15-HETrE which are generated by 12/15-LOX but also by platelet 12-LOX and COXs, were also highly increased and were reduced 50% in *Alox15^-/-^* wounds (Figure 3 B). All these peaked at day 1, then declined subsequently, paralleling the early transient expression of 12/15-LOX in the wound. This indicates that free oxylipin generation from 12/15-LOX is acute and transient, peaking during the inflammatory phase, with lipids being reduced back towards basal levels during the healing phase.

**Figure 3.**
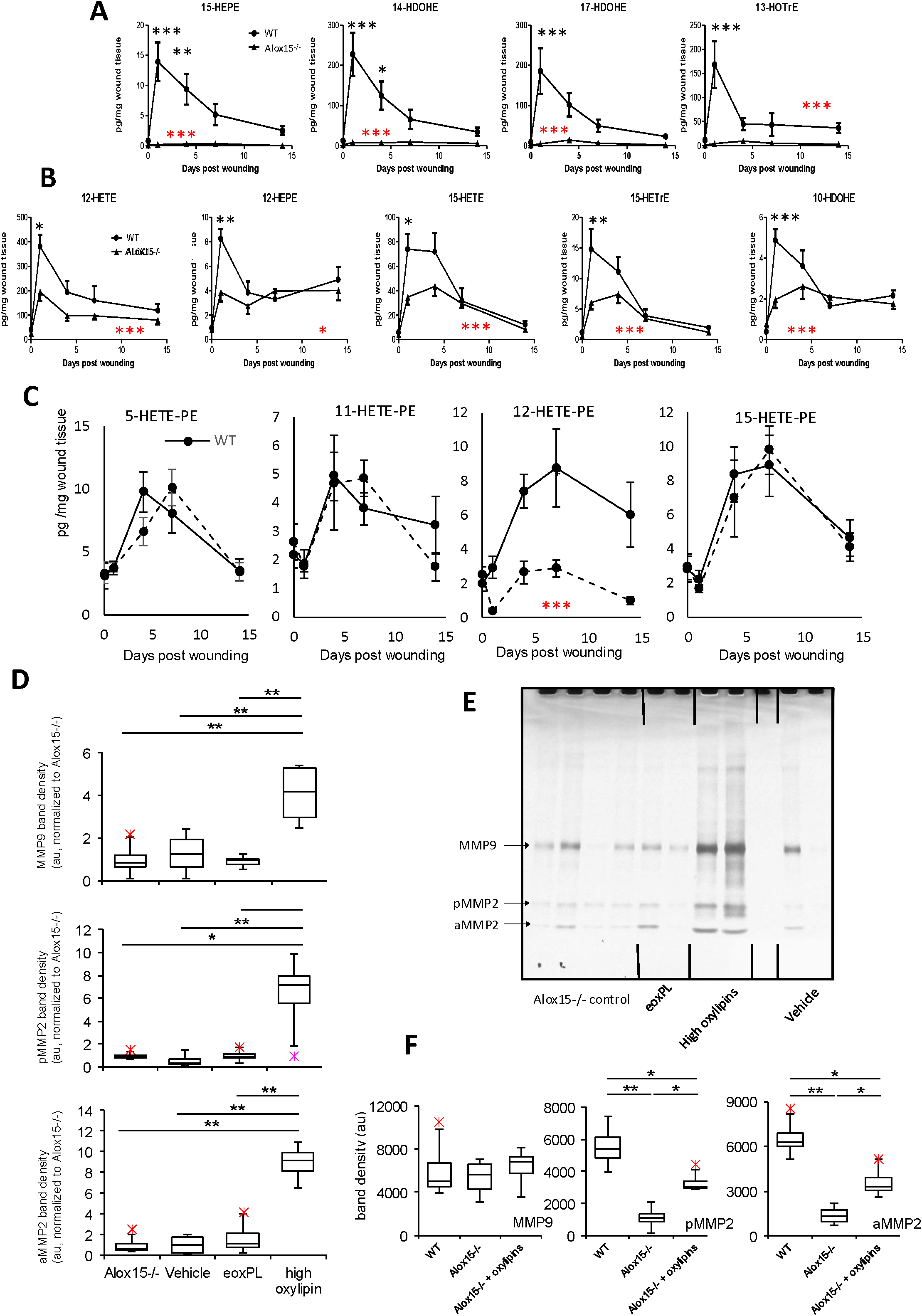
*Alox15^-/-^* wounds show lower levels of many oxylipins and 12-HETE-PEs, while physiological levels of “high oxylipins” restore MMP activity. *Panels A,B. Oxylipins are rapidly elevated post-wounding, but many are reduced in Alox15^-/-^ wounds.* Oxylipins were measured using LC/MS/MS as outlined in Methods. n = 6 samples/time point, with 4 wounds pooled/sample. For all panels differences between groups were analyzed using two-way Anova (red stars), with Bonferroni post hoc test between individual time points (black stars), mean ± SEM, * p < 0.05, ** p<0.01, *** p < 0.005. *Panel C. HETE-PEs elevate during healing, peaking on day 7, with significant loss of 12-HETE-PE isomers in Alox15^-/-^ wounds.* Oxidized phospholipids were measured using LC/MS/MS as outlined in Methods (n = 5 samples/time point, with 4 wounds pooled/sample). Unpaired t-test, * p <0.05, ** p < 0.01, *** p < 0.005. *Panel D. “High oxylipins” restored MMP activities in wounds from Alox15^-/-^ mice*. Post wounding, lipids were added to wounds as indicated in Methods every second day. Wounds were harvested and analyzed for MMP activities using zymography as in Methods (n = 4, 4 – 5 wounds pooled/sample). *Panel E. A gel showing representative data from Panel C*. Panel F. “High oxylipins” restore MMP activities to wild-type level. Wounds were harvested and analyzed for MMP activities using zymography as in Methods (n = 4, 4 – 5 wounds pooled/sample). For Panels C,E, ANOVA with Tukey post hoc test was used, * p <0.05, ** p < 0.01, *** p < 0.005.

Several prostaglandin dehydrogenase (PGDH) metabolites generated from the oxidation of HODEs and HETEs (9-, 13-oxoODE, and 15-oxo-ETE) were similar in both strains, but peaked at day 4 with higher levels in *Alox15*^-/-^ (Supplementary Figure 3 B). Lipids from 5- LOX (5-HETE, LTB4) and COX (PGE2, PGD2, 11-HETE, 11-HEPE, TXB2 and other PGE2 isomers) were strongly elevated at day 1 and declined after but were not impacted by *Alox15*^-/-^ (Supplementary Figure 3 B). The LA products 9-and 13-HODE were elevated early and were slightly lower on day 1 in *Alox15*^-/-^ (Supplementary Figure 3). Several cytochromeP450/soluble epoxide hydrolase (sEH) metabolites were detected at very low amounts with small increases around day 4 which fell by days 7 and 14 (5,6-diHETrE, 8,9-diHETrE, 11,12-diHETrE, 14,15-diHETrE, LTB4, 5,6-EET, 7,8-EpDPA, 13,14-EpDPA) (Supplementary Figure 4). Last, 9,10-EpOME and 12,13-EpOME elevated beyond day 4, although levels fluctuated significantly, similar to their sEH metabolites 9,10-diHOME and 12,13-diHOME (Supplementary Figure 4). None were reduced by *Alox15*^-/-^ indicating they originated from other biochemical or non-enzymatic pathways. Indeed, many from CYP/sEH were significantly higher at day 4. In relation to SPM, out of several monitored, only trace amounts of resolvinD5 were detected, with detailed structural analysis including chiral chromatography shown in Supplementary Methods and Data.

### Enzymatically-oxidized phospholipids (eoxPL) are generated during wound healing, peaking during the proliferative stage, with 12-HETE-PEs significantly impacted by Alox15 deficiency

Free oxylipins generated by COXs or LOXs are also formed as complex lipids attached to membrane phospholipids (PL), termed eoxPL. The most abundant are HETE-containing phosphatidylethanolamine (PE), either generated by direct attack on PE by 12/15-LOX, or by esterification of newly formed HETEs to lysoPE(10). 12/15-LOX generates 12-HETE-containing eoxPL, while 5-HETE-PE arise via 5-LOX (neutrophils)(8, 42). Platelet 12-LOX is a source of 12-HETE-PEs(43). 15-HETE-PEs form either via 12/15-LOX, or through esterification of 15-HETE by COX, which is also a source of 11-HETE-PEs. Following wounding, sustained elevations of 5, 11, 12, and 15-HETE-PEs occurred, peaking on day 4, then declining (Figure 6 A, Supplementary Figure 5 A). Each represents a series of isomers differing by *sn1* fatty acid, with 3-4 per HETE isomer. (Figure 3 C, Supplementary Figure 5 A). 8-HETE-PE were below LOQ indicating that there is little/no non-enzymatic oxidation and confirming that the others are from LOX and COX. Overall, esterified HETEs were less abundant than their corresponding free acid species. Consistent with generation by 12/15-LOX, 12-HETE-PE was reduced by >50% in *Alox15*^-/-^ wounds, while others were unaffected (Figure 3 C). Individual 12-HETE-PE isomers were also all significantly lower in *Alox15*^-/-^ wounds (Supplementary Figure 5 B). These data confirm enzymatic origin, but similar to free HETE, a significant amount is from other sources, for example platelet 12*S*- or skin 12*R*-LOXs. All HETE-PEs peaked around days 5-10, later than free acid HETEs (Figure 3 C). Thus, as free HETEs declined, the esterified forms were elevating. This may reflect onset of esterification processes driven by Lands cycle.

### MMP activities and collagen deposition are restored to wild-type levels in Alox15^-/-^ wounds by high abundance oxylipins, but not eoxPL

Oxylipins are not generated in isolation but as complex mixtures *in vivo*. Here, lipidomics data informed formulation of relevant mixtures to add to healing *Alox15^-/-^* wounds (Supplementary Table 1). *High oxylipins* contained lipids generated acutely in higher amounts that were relatively deficient in *Alox15*^-/-^ wounds, with amounts added via topical dermal delivery aiming to match the maximum levels detected post-wounding (12-HETE, 17-HDOHE, 13-HOTrE, 15-HETE, 12-HEPE, 12-oxo-ETE, 15-HEPE, 12(13)-EpOME, 13-HODE, 14-HDOHE). Wounds treated with *high oxylipins* demonstrated increased levels of MMP9, aMMP2 and pMMP2 (Figure 3 D,E, Supplementary Figure 6 A). Significantly, this treatment reversed the phenotype so that MMP activities were significantly elevated to between WT and *Alox15^-/-^* levels (Figure 3 F, Supplementary Figure 6 B). No impact was seen with a pharmacological dose of PE 18:0a/12-HETE, an eoxPL which was 50 % reduced by *Alox15* deletion (Figure 3 D,E, Supplementary Figure 6 A). Next, the ability of lipids to reduce the accelerated collagen deposition of *Alox15*^-/-^ was tested. Here, treatment with vehicle alone caused a non-significant increase in collagen, but this completely suppressed by the *high oxylipin* preparation, but as for MMPs, there was no impact of eoxPL (Figure 4 A,B).

**Figure 4.**
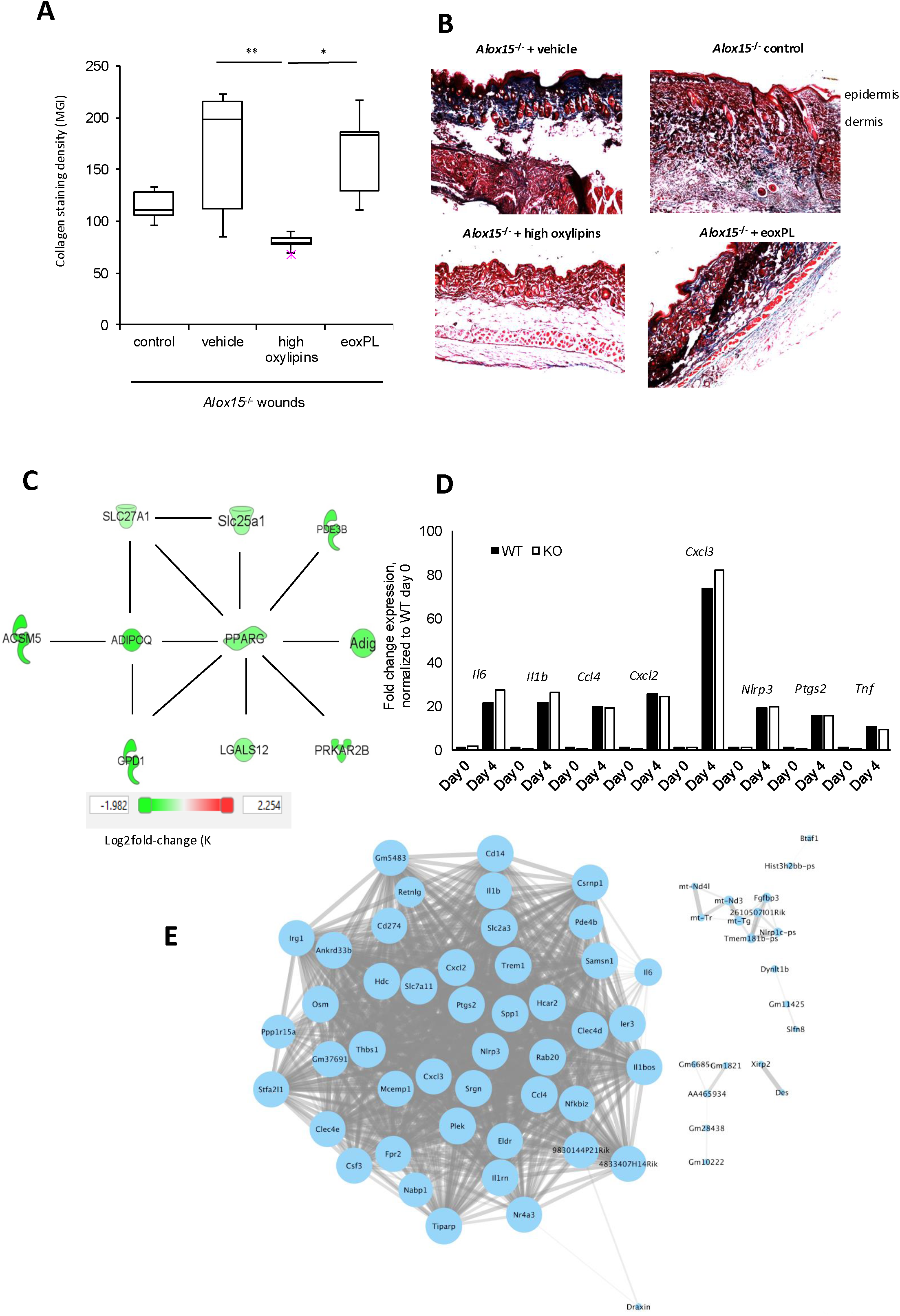
“High oxylipins” dampen collagen deposition, while unwounded *Alox15^-/-^*skin shows reduced PPARγ activity but a normal pro-inflammatory response to wounding, while a series of highly correlated genes fail to revert to baseline in *Alox1*^5-/-^ wounds at Day 7. *Panel A. Collagen generation is dampened by “high oxylipins”.* Post wounding, lipids were added to wounds as indicated in Methods every second day. Wounds were harvested at Day 7 and analyzed for collagen using Masson’s Trichrome staining (collagen: blue, epithelial cells: deep red, nuclei: black, non-collagen structures: pink) and pixels counted (n = 5-6 wounds/group. *Panel B. Representative images from Panel A. Panel C. Unwounded Alox15^-/-^ skin shows significantly reduced PPAR*γ*/adiponectin expression/activity.* RNASeq was carried out as indicated in Methods on non-wounded skin (n = 3 – 4 samples per group, each sample was a pool of 4 wounds per animal with 2 animals per pool = 8 wounds per sample). All genes shown are significantly reduced in *Alox15^-/-^* skin, and are either controlled by or regulate PPARγ/adiponectin. *Panel D. Upregulation of a series of canonical “inflammatory” genes is preserved in Alox15^-/-^ wounds.* Gene expression was normalized for each gene to its Day 0 mean value, then expressed as fold-change (n = 3 – 4 per group) each sample was a pool of 4 wounds per animal with 2 animals per pool = 8 wounds per sample). For Panel A, ANOVA with Tukey post hoc text. *Panel E. A large number of genes that are significantly different between wild-type and Alox15^-/-^ wounds at Day 7 highly correlate across the whole timecourse, indicating coordinated regulation*. Genes that were found to be significantly differentially expressed at Day 7 were analyzed in Cytoscape, using their expression levels for the entire time-course, with correlation [r] > 0.8 shown.

### Transcriptional analysis reveals reduced anti-inflammatory lipid metabolism in Alox15^-/-^ healthy skin, but a relatively normal acute response to wounding

To identify transcriptional networks modulated by *Alox15^-/-^,* RNASeq was performed on day 0 (healthy tissue), and days 4 and 7 post-wounding. At Day 0, 143 genes were significantly different between the two strains (adjusted p-value < 0.05) (Supplementary Table 6). Analysis of these using Ingenuity Pathway Analysis (IPA) identified lipid metabolism as highly represented, and a subset of relevant genes in that network is shown (Figure 4 C). Strong downregulation of *Adipoq* (adiponectin) and *Pparg* (PPARγ) was seen, associated with a reduction in a series of genes that either control or are controlled by these (Figure 4 C) (44–55). PPARγ is a transcription factor that induces adiponectin(56), and it responds directly to oxylipin ligands generated by *Alox15* including several HODEs, HETEs and HDOHEs(13, 16–19). Adiponectin is a hormone and adipokine centrally involved in metabolism, that is protective against a number of inflammatory conditions such as atherosclerosis and type 2 diabetes(57, 58). Both are crucial in mediating anti-inflammatory actions such as inhibition of pro-inflammatory NFkB signaling and NLRP3(59, 60). Additional down-regulated genes in the network include regulators of lipid metabolism such as *Slc27a1* (import of long-chain fatty acids), *Acsm5* (Acyl-CoA synthetase medium-chain family member 5), *Gpd1* (regulates lipid metabolism), and two genes that regulate adipose tissue development, *Adig* (adipogenin) and *Lgals12*. Importantly, reduced basal expression of *Adipoq* and *Pparg* indicates that *Alox15^-/-^*tissues would struggle to mount the anti-inflammatory response required to counterbalance inflammation during the later wound healing phase in which PPARγ signaling is known to play a role(61). At day 4, comparison between WT and *Alox15^-/-^* wounds showed that around 42 genes were significantly different (Supplementary Table 7). These genes didn’t appear obviously functionally related. However, the classic inflammatory response to injury, as measured by genes that include *Tnfa, Il1b, IFNg, Nlrp3, Cxcl2, Ccl4*, and *Il6* was preserved in both strains indicating a normal onset of inflammation (Figure 4 D).

### Late wound healing in Alox15^-/-^ wounds shows a failure of inflammation to reduce to basal levels, with many pro-inflammatory genes remaining upregulated

At day 7, 79 genes were significantly different between the strains, with 60 higher in *Alox15*^-/-^ wounds than WT (Supplementary Table 8). A Cytoscape analysis was performed using the whole-time course dataset for the 79 genes (day 0, 4, 7 and both strains), and a sub-group of 45 were seen to strongly correlate, suggesting their behavior was co-ordinated during the entire wounding and healing process (Figure 4 E). These included several pro-inflammatory genes such as *Ccl4, Cd14, Cd274, Clec4d, Clec4e, Csf3, Cxcl2,Cxcl3, Fpr2, Il1b, Il6, Irg1, Nfkbiz, Nlrp3, Ptgs2, Retnlg, Trem1* and *Osm*. Many are involved in macrophage-driven inflammation, and all were highly upregulated in both WT and *Alox15^-/-^* wounds on day 4. However, in *Alox15^-/-^* wounds these all failed to reduce back to basal levels at day 7, with their transcription remaining around 50% of the day 4 levels (Figure 5 A, Supplementary Figure 7). This contrasts with WT wounds where these fully returned to basal levels by day 7 (Figure 5 A, Supplementary Figure 7). IPA analysis of this sub-group of genes demonstrated that many are upregulated through common mechanisms, such as NFκB. For example, within this group, an IPA sub-network predicted higher activity of the IL1, IFNβ and Inflammasome pathways in the *Alox15^-/-^* wounds. Importantly, IL1, IFNβ and Inflammasome are all well known to be downregulated by PPARγ (Figure 5 B). Indeed, several genes in this network are also known to be down-regulated by activation/induction of PPARγ either directly or via inhibition of NF-κb, including *Il6, Nlrp3* and *Il1b* (60, 62, 63). Last, it was seen that while *Pparg* expression was not induced by wounding, its expression fell in WT wounds to levels similar to those seen in *Alox15^-/-^* (Figure 5 C). This suggests that the lack of PPARγ signalling during the healing response results from a relative deficiency in 12/15-LOX- derived ligands, explaining why their supplementation could reduce the fibrotic phenotype in *Alox15^-/-^* wounds.

**Figure 5.**
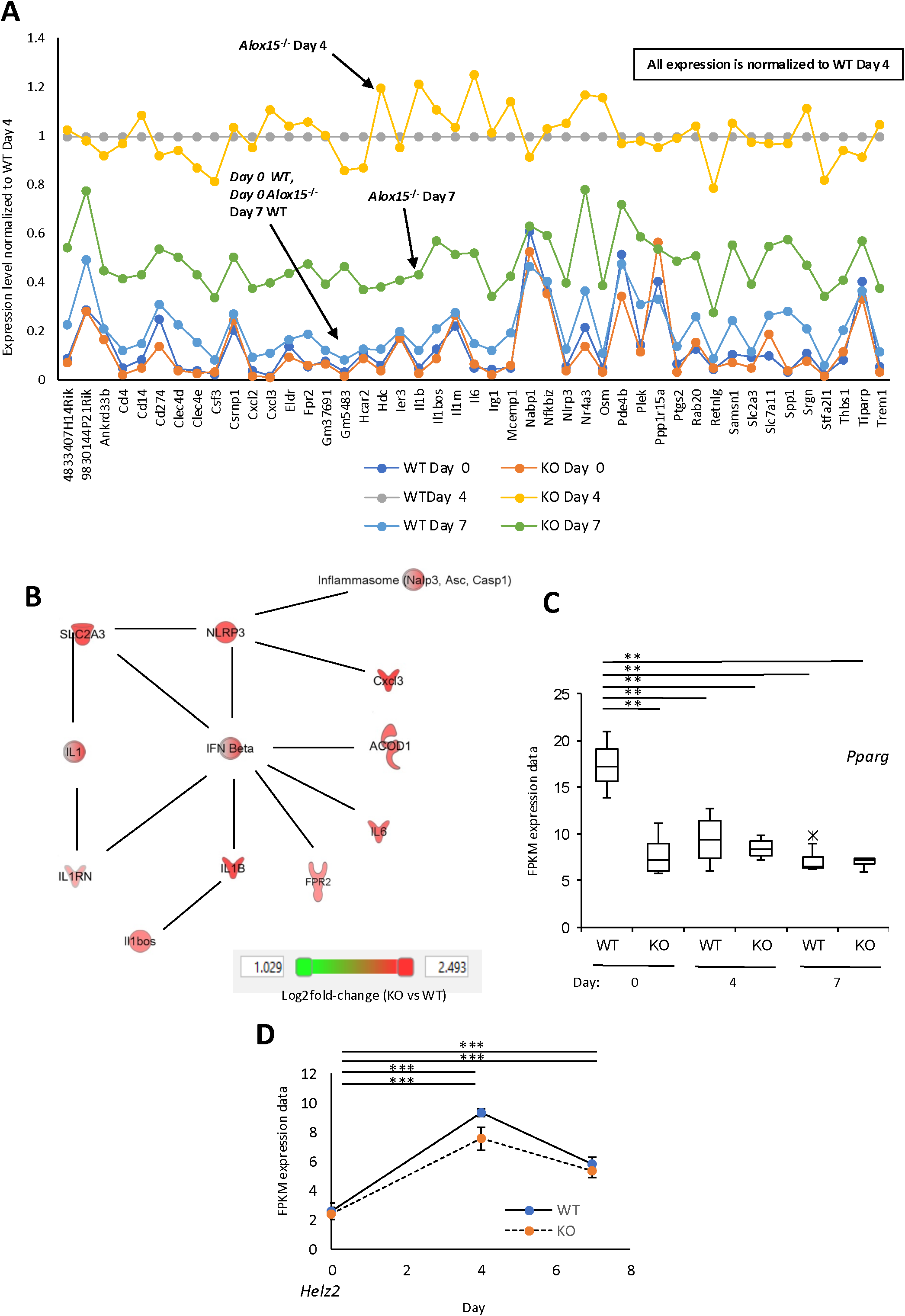
Genes that fail to revert at day 7 in *Alox15^-/-^* wounds remain 50% elevated above baseline, and many are controlled through NLRP3, IFNβ and IL-1, PPARγ expression is not upregulated during wounding, and *Elf4 and Helz2* are upregulated during wounding. *Panel A. All genes from the highly correlated network in* Figure 4 *E typically remain 50% elevated.* Data from gene expression of highly correlated genes was averaged and normalized to day 4, wild-type mean (the inflammatory response level) (n = 3 – 4 per group). *Panel B. IPA network analysis of genes that are significantly different at Day 7 reveals significantly higher levels of Inflammasome, IFNb and IL-1 pathways*. Gene expression data from Day 7 was analyzed using IPA. *Panel C. Pparg expression is not increased during wounding*. Transcriptional data on *Pparg* was compared across the timecourse (n = 3 – 4 per group). For all gene expression data, students t-test, followed by Benjamin Hochberg correction: * p <0.05, ** p < 0.01, *** p < 0.005. *Panel D.* Data from gene expression is shown for WT and Alox*15^-/-^* wounds during the time course (n = 3 – 4 per group). For all gene expression data, students t-test, followed by Benjamin Hochberg correction: * p <0.05, ** p < 0.01, *** p < 0.005.

“High oxylipins” generated during the wound response induce transcriptional activity of PPARγ in a reporter assay.

To determine whether the more abundant oxylipins generated by 12/15-LOX during wounding activate PPARγ, the *high oxylipin* mixture was tested in a reporter assay, with HEK293 cells expressing mouse PPARγ and *Firefly* luciferase under control of 3x Ppar Responsive Element (PPRE)(40) exposed to the lipids for 24 hrs. Doses tested were informed by oxylipin amounts detected in wounds in this study and studies which show *in vitro* oxylipin activation of PPARγ in the 10-100 μM range(13, 15–18). First, amounts detected in an individual wound on Day 1 were tested (Supplementary Table 1). These were diluted into 50 μl media giving a final concentration of 0.4 – 1.2 μM total oxylipins. However, at these doses, PPARγ was not reliably activated (not shown). Thus, we next tested amounts of oxylipins previously shown to bind and activate PPARγ *in vitro*. The most abundant lipid in our mixture was 12-HETE, which ranged from 12-36 μM at the three doses tested, with total oxylipins at 40-120 μM (Supplementary Table 1). Activation of PPARγ was seen for all oxylipin doses, with 80 μM showing a significant increase (Supplementary Figure 8 A). These data are in line with previous reports that many individual 12/15-LOX products can act as low-affinity PPARγ ligands at these concentrations, and that *Alox15*-deficiency leads to loss of PPARγ activation in macrophages(13, 15–19). Overall, our data shows that the mixture of oxylipins can act activate PPARγ in a complex mixture, in full agreement with previous literature.

### Wounding induces significant increases in gene expression of the PPARγ co-activator Pdip1/Helz2/Pric285

We noted that the amounts of oxylipins used *in vitro* were higher than we detected *in vivo*. However, exact wound oxylipin concentrations are not possible to determine, and it is also not known how much oxylipin enters the cells to bind and activate PPARγ *in vitro*. Furthermore, the ability of oxylipins to activate PPARγ may be regulated by known co-activators present *in vivo*, including free fatty acids and protein co-activators(64, 65). Prompted by this, the expression of known protein co-activators was next interrogated in transcriptional data(65). One showed consistent and significant upregulation in both WT and *Alox15^-/-^*wounds on both days 4 and 7, versus day 0 (Figure 5 D), with highest expression seen on day 4. This protein, PPARγ-DBD-interacting protein 1a, HELZ2/PRIC285 (*Helz2*) is a helicase that binds DNA binding domains of PPARγ through its C-terminal region, and can enhance PPARγ activation by troglitazone directly(66). Whether HELZ2/PRIC285 can similarly enhance the ability of oxylipins to bind and activate PPARγ is unknown and remains to be tested.

### Comparison of temporal changes in gene expression indicates additional transcription activators regulated by Alox15 beyond PPARγ, including Elf4, Cebpb and Tcf3

Last, RNASeq was deeper interrogated for additional *Alox15*-modulated transcriptional regulators, using a temporal analysis. Here, we characterized individual strains separately and found three new candidates (see Supplementary Methods and Data for full analysis). *Elf4* is a known anti-inflammatory transcription regulator of inflammation, which targets several genes in the list, including *Anln, Asf1b, Ccnb2, Cdca3, Cenpa, Cenpe, Cks2, E2f8, Hmmr, Kif4a, Mcm10, Ndc80, Oip5, Rrm2, Tpx2*(*67*). *Tcf3* promotes cell migration and wound repair (68), and *Cebpb* is involved in macrophage repair responses and inflammation (69, 70). Many genes mapping to networks that regulate cytokines were also identified, further evidencing the impact of *Alox15* on inflammatory signaling and identifying a large number of novel targets for further study (see Supplementary Methods and Data).

## Discussion

In this study, lipidomic and transcriptomic analyses of *Alox15^-/-^* mice together reveal significant pathways which are impacted by deletion of the enzyme during wound healing. Overall, many pro-inflammatory genes highly induced by wounding fail to return to basal expression in *Alox15^-/-^* wounds. Taken with our phenotypic data and the response to known PPARγ ligands from this pathway, an intrinsic anti-inflammatory action of 12/15-LOX, driven by the PPARγ/adiponectin axis is lost in *Alox15^-/-^* mice. This allows an uncontrolled fibrosis response, which is almost identical to that previously reported in PPARγ-deficient mice (described below)(71). Transcription factors, including *Elf4*, *Tcf3* and *Cebpb* may also play important roles, but their precise functions remain to be established.

12/15-LOX generates abundant mono-oxygenated oxylipins, eoxPL and in concert with other LOXs is proposed to generate rarer multiply oxygenated SPM via transcellular biosynthesis. However, which lipids are primarily responsible for the effects of 12/15-LOX during physiological healing/resolution have been unclear. Its lipids are not generated in isolation but in complex mixtures of varying abundance but studies testing their role in inflammation have usually added them singly(21–24). To test the role of the enzyme under physiological conditions, we adopted a well-characterized model of wounding that fully heals within 14-days. Transcriptomic, phenotypic and lipidomic approaches revealed that *Alox15* deletion leads to an accelerated healing response resulting in a “fibrotic” phenotype. During this, higher collagen deposition, increased stem cell proliferation and differentiation are seen, as well as higher levels of inflammatory markers such as IL6/pSTAT3, pSMAD3 and IFNγ. A failure of pro-inflammatory gene expression to reduce back to baseline during the later remodeling phase was found. Our transcriptional data identified a basal deficiency of PPARγ expression and activity and also showed that many genes which are upregulated by transcription factors such as NLRP3 or NFkB (both antagonized by PPARγ) did not revert to basal levels post-wounding. Linked with this, wild-type wounds generated large amounts of several known PPARγ ligands via 12/15-LOX (e.g. 12-HETE, 15-HETE, 12-HEPE, 13-HOTrE) during the early inflammatory phase, and critical features of normal wound healing could be restored in *Alox15^-/-^* by supplementing with physiological levels of a mixture of these(13, 15–20). Furthermore, several oxylipins that were deficient in *Alox15^-/-^* wounds are known to dampen IL-6 (12-HETE, 15-HETE, 14-HDOHE, 15-HEPE(7, 15, 72)) and NLRP3 (13-HOTrE(73)), with these effects also likely mediated by PPARγ. In vitro testing established that the mixture of oxylipins could activate PPARγ *in vitro*. A caveat is that concentrations needed, although in line with many other studies on PPARγ, appeared to be around 100-fold higher than those found *in vivo*. This suggests that the lipids alone are not sufficient and protein co-activators known to sensitize PPARγ to agonists may be involved, such as HELZ2/PRIC285 (*Helz2*) (*66*), which we also found to be upregulated significantly during the wounding response. Based on our staining data, the most likely cellular sources of the lipids will be F480^+ve^ macrophages and stem cells at the base of hair follicles.

Notably, the phenotype seen in our study is almost identical to that of PPARγ-deficient mice, where increased actin, collagen pSMAD3 and an accelerated healing/fibrotic phenotype in skin were described(71). Also, loss of PPARγ in skin fibroblasts is associated with elevated pSMAD3, while PPARγ agonists directly reduce actin and collagen expression(74, 75). Furthermore, PPARγ blockade elevates MMP1 and MMP9 in fibroblast-like synoviocytes(76) while its activation dampens fibroblast proliferation and differentiation(77). PPARγ also inhibits expression of IL6, IFNγ and pSTAT3, and prevents pSMAD3 dependent collagen synthesis and deposition in fibroblasts(78–81). Thus overall, the *Alox15^-/-^* fibrotic phenotype most likely results from a simple failure to generate mixtures of abundant PPARγ ligands during the acute response to injury. In line with this, an anti-inflammatory and pro-healing action of pharmacological PPARγ agonists such as rosiglitazone in mouse wound models of diabetes and obesity has been described previously(82, 83).

As well as free oxylipins, eoxPL were detected, elevating significantly during the remodeling stage, with 12-HETE-containing isomers reduced almost to basal levels in *Alox15^-/-^* wounds. eoxPL are also known as PPARγ ligands, although added herein, they did not restore MMP/collagen, most likely due to insufficient concentration (84, 85). Apart from very low amounts of RvD5, SPM were not generally detected. A recent study on murine cutaneous wounds reported several RvDs in mouse wounds, including RvDs1,2,3,4,5,6 and two 17*R*- isomers and proposed a central role for these lipids in repair through reconstitution studies with exogenous lipids(86). There, administration of individual RvDs at around 100 ng/wound/day per isomer demonstrated a pharmacological effect on healing(86). Oxylipin amounts administered in our study (6.5 ng total dose/wound/day) were considerably lower. Thus, in the previous study, the mechanism could involve SPM acting *via* low affinity PPARγ binding and activation, similar to other oxylipins. In this regard, the closely related RvD1 (7*S*,8*R*,17*S*-triHDOHE) was previously reported to activate PPARγ in a mouse model of acute lung injury(87). Alternatively, a recent study showed that RvDs can allosterically activate the PGE_2_ receptor, EP4, with RvD5 sensitizing at nM concentrations (34). In our study, 7,17-diHDOHE (RvD5) was detected at extremely low amounts (max amount 1.5 pg/wound, equating to around 4 fmol/wound). Although it’s not possible to calculate local concentrations in wounds, these amounts appear too low to mediate EP4 sensitization or PPARγ activation. In our study, the doses of oxylipins that were bioactive *in vivo* appeared to be around 100-fold lower than required for PPARγ activation *in vitro*. The upregulation of co-activators such as *Helz2* may provide a partial explanation since it can sensitize PPARγ to agonists(66). This remains to be experimentally tested in relation to oxylipins.

PPARγ binds and is activated by many diverse lipid ligands with relatively low affinity and little differentiation of enantiomeric structure. Thus, the concerted action of many agonists generated in relatively high amounts during the healing process is consistent with the known role of PPARγ in mouse wound healing(71). Here, our studies using *Alox15^-/-^* mice support the idea that abundant lipids generated by the 12/15-LOX pathway act in concert to promote the well-known anti-inflammatory actions of this transcription factor, preventing uncontrolled fibrosis through dampening inflammation directly. This is in line with the long-known action of many 12/15-LOX monohydroxy ligands as PPARγ ligands, and previous reports of defective PPARγ signaling in *Alox15^-/-^*macrophages(13), and supports consideration of therapies targeting this pathway in defective wound healing in patients.

## Acknowledgements

Funding is acknowledged to VOD and CPT from the Medical Research Council (MR/M011445/1), and from European Research Council (LipidArrays). We acknowledge Sarah Edkins and Shelley Rundell for technical support. Wales Gene Park is a Health and Care Research Wales-funded infrastructure support group. VOD acknowledges the Royal Society Wolfson Merit Award Scheme. Funding to AVP is acknowledged by the European Research Council (ERC, grant agreement No 669879). MP was funded by Wellcome Trust GW4-CAT Fellowship (216278/Z/19/Z). SC acknowledges a grant PID2021-125406B-100 from MCIN/AEI /10.13039/501100011033/ and by ERDF a way of making Europe. We gratefully thank Hartmut Kühn, Humboldt University Berlin for the gift of antibody directed against 12/15-LOX, and Paul Martin, University of Bristol for assistance with the murine wound model.

## Authorship contributions

JJB, SRCJ, VJT, MA, JAJ, RI, AL, LF, JC, BCC, CG, AC, SC conducted experiments. JJB, CPT, VBO designed experiments. JJB, VBO, CPT, AJC supervised experiments. SAJ, JJB, VBO, CPT, AVP, SC interpreted findings and provided additional intellectual input. JJB, VBO drafted the manuscript. RA, BS performed statistical and informatic analysis of RNASeq data. All authors edited the manuscript.

## Supplementary Methods and Data

### Supplementary Methods

#### Animal model

Mice (8-12 weeks old C57/B6/J) were purchased from Charles River UK (Margate, UK), while *Alox15^-/-^* mice were bred in house (F11, C57BL/6J) in isolators. All animal experiments were performed in accordance with the United Kingdom Home Office Animals (Scientific Procedures) Act of 1986, under License (PPL 30/3334). Male and female mice were used for all studies except RNASeq where only males were used to reduce variation. Mice were housed in scantainers on a 12-hour light/dark cycle at 20 - 22 °C, with free access to regular chow and water. Mice were anaesthetized using 3 - 3.5 % isoflurane delivered in 2 L per min 100 % oxygen. Once areflexic, mice received a sub-cutaneous dorsal injection of 10 μl Temgesic/Buprenorphine (1 μg). They were shaved and re-tested to confirm the areflexic state. Spinal midline was drawn before rotating to one side. Skin was folded using the spinal midline and two punch biopsy needles (BD pharma, UK) used to create 4 wounds (2 wounds per 4 mm biopsy needle)(1). Wounds were trimmed using clean scissors, and wounds photographed for size. Mice were transferred to a warming box until regaining consciousness, before transfer into cages lined with paper towels. Mice were monitored at 1 - 3 hours post wounding, and before the end of the light cycle. At 24 hours, mice were transferred into their original cages. In some experiments, mixtures of lipids were applied to wounds to determine their impact on healing. Two preparations were used, either “high-abundance” or “oxPL”. Here, immediately post-wounding, lipids were added (as described in Supplementary Table 1) in a final volume of either 50 μl (days 0,2) or 25 μl (days 4,6), with amount of lipids consistent across all days. Lipids were added in ethanol:Tween80:sterile water (1:1:18) as vehicle(2–4). On days 2, 4, 6, mice were briefly anesthetized as above to enable lipid application. Vehicle or lipids were added topically to the wound and allowed to sit in the wound surface for 15 min, before mice were allowed to recover consciousness. On various days post wounding, mice were euthanized and wounds collected (typical weights from 5-10 mg per wound) and either processed for histology as outlined below using paraformaldehyde fixation or snap frozen in liquid nitrogen and then stored at -80 ^0^C. For lipid supplementation studies, mice were euthanized at day 7.

#### Generation of histological tissue sections

At various time points up to 14 days, mice were killed using CO_2_ (Schedule 1). The wound site and a small area of surrounding tissue was dissected and placed in 4 % paraformaldehyde for 72 hours, then 70% ethanol (to prevent further crosslinking), then the tissue processed for paraffin wax embedding. Wounds were placed into plastic cassettes and in a Leica tissue processor using the following parameters: ethanol (60 %) under ambient temperature under vacuum for 90 min with agitation. Ethanol (70 %) under ambient temperature under vacuum for 90 min with agitation. Six separate steps of ethanol (100 %) under ambient temperature under vacuum for 60 min with agitation. Xylene (100 %) at 37 °C under vacuum for 120 min with agitation. Xylene (100 %) at 45 °C under vacuum for 120 min with agitation. Two separate steps of wax (100 %) at 60 °C under vacuum for 120 min with agitation. Wax (100 %) at 60 °C under vacuum for 60 min with agitation. Wax (100 %) at 60 °C under vacuum for 45 min with agitation. Post-processing, plastic cassettes were removed, and wounds were cast side on in molten (60 °C) wax, which was allowed to harden to form tissue wax blocks. Wax blocks were fastened to a microtome stage and secured. 10 μm slices were taken from each revolution and transferred to a 40 °C water bath and allowed to unfold and float for 30 - 60 seconds before being placed onto a glass slide. Sections were left to dry at room temperature, then at 55 °C overnight.

#### DAB Immunohistochemistry (F480, LY6G)

Slides were placed in Histoclear (Fisher Scientific) for 3 min at room temperature then into ethanol at 100 %, 90 % and 70 % for 3 min each before being placed into running tap water for 5 min. Next, slides were dried with a paper towel, and a hydrophobic pen used to draw borders around each section. Antigen retrieval was performed by addition of 5μg/ml proteinase K in PBS for 7 mins at R.T. Slides were then rinsed in running tap water and H_2_O_2_ (3 %) was added and the sections incubated for 10 min at room temperature. Slides were washed in running tap water. Block solution (500 µl PBS + 1 % bovine serum albumin + 0.1 % fish gelatin + 0.5% Triton X-100) and avidin block (Avidin/Biotin block kit vector labs) were added and sections incubated for 30 min at room temperature. Block solution was removed, and slides washed twice with PBS. Primary antibody (Supplementary Table 2) or isotype controls made up in antibody solution (50 ml PBS + 1 % bovine serum albumin + 0.1% fish gelatin + 0.5% Triton X-100 and biotin block solution) was then added F4/80 Ab at 1/400 and Ly6G at 1/200. Slides with antibody solution were left for an extended period (14 hours or overnight) then washed with PBS 0.1% Tween 2 times for 5 mins. Biotin conjugated secondary antibody solutions were made up at 1:500 in block solution and added after the washes for 30mins. Secondary antibody was removed and sections washed 2 times as done before. Avidin-Biotin Complex (ABC) kit solution (Vector Labs) was then made up (as per manufacturers’ instructions) 30 min before use, this was added for 30 min before being removed with one 5 minute wash. Sections were treated with a 3,3’Diaminobenzidine (DAB) solution (Vector Labs) for 1-2 min depending on staining intensity. The sections were washed in running tap water for 5 mins, then placed in hematoxylin for 15 secs. Sections were washed for 5 min in running tap water. Slides were then dipped 3 times in acid alcohol and then washed in running tap water for 1 min. Finally slides were placed in Scotts Tap water for 18 seconds, followed by a final wash in running tap water for 5 mins. Slides were then placed into increasing concentrations of ethanol (70 %, 90 %, 100 %) for 30 secs each, then placed into Histoclear for 2x 30 sec incubations. Sections were removed from Histoclear and dried carefully using paper towels. Slides were treated with 2 drops (50 μl) of distyrene and xylene (DPX solution) and covered with a glass cover slip and put in an oven at 60°C overnight.

#### DAB Immunohistochemistry (all other antigens)

Slides were placed in Histoclear (Fisher Scientific) for 2 min at room temperature then into ethanol at 100 %, 90 % and 70 % for 5 min each before being placed into DDH_2_O for 5 min. Slides were then placed in citrate buffer (10 mM sodium citrate, 0.05 % Tween 20, pH 6.0) and incubated at 96 °C for 1 hr. They were cooled to room temperature over 30 min, then placed back into DDH_2_O for 30 min. Next, slides were dried with a paper towel, and a hydrophobic pen used to draw borders around each section. Phosphate buffered saline (PBS, 1 ml) was added to sections for 2 min before removal using a Pasteur pipette. H_2_O_2_ (3 %) was added and the sections incubated for 10 min at room temperature. Slides were washed twice using PBS. Block solution (500 µl PBS + 1 % bovine serum albumin + 0.1 % fish gelatin) and avidin block (Avidin/Biotin block kit vector labs) was added and sections incubated for 45 min at room temperature. Block solution was removed, and slides washed twice with PBS. Primary antibody (Supplementary Table 2) or isotype controls made up in antibody solution (50 ml PBS + 1 % bovine serum albumin + 0.1% fish gelatin and biotin block solution) was then added. Slides with antibody solution were left for an extended period (14 hours or overnight) then washed with PBS 5 times with 10-minute section submersion in between washes. Secondary (HRP conjugated) antibody solutions were made up at desired concentration (Supplementary Table 2) in block solution and added after the washes for two hours. Secondary antibody was removed and sections washed 5 times as done before with 10-minute gaps between wash steps, Avidin-Biotin Complex (ABC) kit solution (Vector Labs) was then made up (as per manufacturers’ instructions) 15 min before use, this was added for 45 min before being removed with 5 washes. Sections were treated with a 3,3’Diaminobenzidine (DAB) solution (Vector Labs) for 5 - 7 min depending on staining intensity. The sections were washed multiple times with DDH_2_O, then placed in hematoxylin for 2.5 minutes. Sections were washed for 5 min in DDH_2_O. then placed into increasing concentrations of ethanol (70 %, 90 %, 100 %) for 2 min each, then placed into xylene for 20 sec. Sections were removed from xylene and dried carefully using paper towels. Slides were treated with 2 drops (50 μl) of distyrene and xylene (DPX solution) and covered with a glass cover slip and stored 24 hrs to dry.

#### Fluorescence Immunohistochemistry

Slides were placed in Histoclear (Fisher Scientific) for 2 min at room temperature then into ethanol at 100 %, 90 % and 70 % made up in DDH_2_0 for 5 min each before being placed into DDH_2_O for 5 min. Slides were placed in citrate buffer (10 mM sodium citrate, 0.05 % Tween 20, pH 6.0) and incubated at 96 °C for 1 hr. They were cooled to room temperature during 30 min, then placed into DDH_2_O for 30 min. Next, slides were dried with a paper towel, and a hydrophobic pen used to draw borders around each section. PBS (1 ml) was added to sections for 2 min before removal using a Pasteur pipette. Block solution (500 µl PBS + 1 % bovine serum albumin + 0.1 % fish gelatin) was added and sections incubated for 45 min at room temperature. Block solution was removed and primary antibody (Supplementary Table 2) or isotype controls made up in antibody solution (50 ml PBS + 1 % bovine serum albumin + 0.1 % fish gelatin) were then added. Slides with antibody solution were left for an extended period (14 hours or overnight). The slides were then washed with PBS 5 times with 10-minute section submersion in between washes. Secondary antibody solutions were made up at 1:300 in block solution and added after the washes and left for 2 hrs at room temperature. Secondary antibody was removed and sections washed 5 times as done before with 10-minute gaps between wash steps. Sections were counterstained with DAPI solution (Invitrogen, 10ng/ml) and True Black (Cambridge BioSciences, Cambridge UK) as per manufacturer’s instructions. Sections were washed 5 times in DDH_2_0 and then allowed to dry overnight and then coverslipped with ProLong™ Diamond Antifade mounting medium (Invitrogen).

#### Collagen staining using Masson trichrome

Collagenous fibres were visualized using Masson trichrome stain (Abcam, ab150686). Paraffin embedded sections were re-hydrated in decreasing concentration of ethanol (100 %, 96 %, 70 % in DDH_2_O, then manufacturer’s instructions followed: placing sections in Bouin’s fluid, Weigert’s iron hematoxylin solution, Biebrich scarlet/acid fuchsin solution, phosphomolybdic/phosphotungstic acid solution, aniline blue solution and acetic acid solution. Sections were dehydrated in increasing concentrations of ethanol (70 %, 96 %, 100 %), before being rinsed in xylene, dried and coverslipped with DPX mounting media. Sections were visualized under a light microscope Leica DM 2000 with a Leica DMC 2900 camera.

#### Image acquisition and analysis

Images were acquired with ether a light microscope (as detailed above) or epifluorescent microscope (EVOS M5000). Images were transferred to ImageJ software for post-acquisition analysis.

i. For DAB images, contrast was enhanced by 5 % before isolation of the brown (DAB) staining channel after a color devolution set to ‘H DDAB’ (hematoxylin and DAB staining). Brown (dab staining) channels were then inverted and set to ‘rainbow smooth’ allowing localization and intensity of the DAB staining to be determined. Pixel count of the image was determined using a live histogram with data only from 70-255 (grey scale) to avoid background (hematoxylin) staining.
ii. For fluorescent imaging pixel count or mean grey intensity (for Texas red or CY5 channels) was used. Isotype controls used are provided in Supplementary Figure 9.

#### RNASeq

Wound tissue dissected from 2 mice (8 wounds in total, 4 wounds per mouse) to generate one sample were snap-frozen in liquid N_2_ before being stored at -80 ^0^C. For WT and *Alox15^-/-^* a total of four samples were generated at each timepoint. Wounds were placed frozen into a pre-cooled pestle and mortar, then ground into a fine powder with liquid N_2_ added. Tri reagent (1 ml, Sigma-Aldrich, UK) was added. The sample was then transferred to RNAse-free tubes and bromochloropropane (200 µl, Sigma-Aldrich, UK) added before vortexing. Samples were placed on ice for 5 min, before being centrifuged at 15,000 g for 15 min at 4 °C. The upper aqueous phase was transferred to a new tube and 250 µl of 3 M sodium acetate (pH 5.5), 700 µl (100 %) propanol and 10 µl glycogen added, before incubation at -80 ^0^C overnight. Samples were centrifuged at 15,000 g for 15 min at 4^0^ C to pellet the RNA. The pellet was washed 3 times using 70 % ethanol, then allowed to air dry for 5 min. RNAse-free water (50 µl) was added. Samples were cleaned using an RNeasy MinElute Cleanup Kit (Catalogue number 74204 Qiagen, MD, USA), using a column based clean up step. Sample (1 µl) was analyzed using a NanoDrop™ 2000/2000c Spectrophotometer (ThermoFisher Scientific, Newport UK), and to ensure samples were free of contamination. All samples had absorbance 260/230 ratios of between 1.7 and 2.0, and 260/280 values between 1.8 to 2.1. Each RNA sample (5ul) was used to determine RNA integrity analysis (RIN). Total RNA quality was assessed using the Agilent 4200 TapeStation with RNA ScreenTape® (Agilent Technologies) and quantity with the Invitrogen™ Qubit-iT™ RNA HS Assay Kit (Fisher Scientific) according to the manufacturer’s instructions. Libraries were prepared from 300ng of total RNA with a RIN value > 7. Total RNA was depleted of ribosomal RNA and sequencing libraries prepared with the Illumina®TruSeq Stranded Total RNA Library Prep Gold (Illumina, Inc) kit using TruSeq CD Index Adapters1 (Illumina, Inc). The steps included the depletion of cytoplasmic and mitochondrial rRNA, depleted RNA fragmentation, 1st strand cDNA synthesis, 2nd strand cDNA synthesis, adenylation of 3’ ends, adapter ligation, DNA fragment enrichment by PCR amplification (10-cycles) and validation. The manufacturer’s instructions were followed except for the cleanup after the ribozero depletion step where Ampure®XP beads (Beckman Coulter) and 80% Ethanol were used. The libraries were validated using the Agilent 4200 TapeStation® with high-sensitivity D1000 ScreenTape® (Agilent Technologies) to ascertain the insert size, and the Invitrogen™ Qubit™ dsDNA HS Assay Kit (Fisher Scientific) used to perform the fluorometric quantitation. Following validation, the libraries were normalized to 4 nM, pooled together and clustered on the cBot™2 (Illumina, Inc) following the manufacturer’s recommendations. The pool was then sequenced using a 75-base paired-end (2x75bp PE) dual index read format on the HiSeq4000 (Illumina, Inc) according to the manufacturer’s instructions. These combinatorial dual (CD) index adapters were formerly called TruSeq HT.

#### Lipid extraction

At time points < 14 days, mice were killed using CO_2_ (Schedule 1). The wound site and a small area of surrounding tissue was dissected. Wounds were placed into 0.3 ml buffer containing 100 μM diethylenetriamine pentaacetate. Ceramic beads (15 – 20, 2.8 mm) were added (approx. 10/sample) and tubes placed into a bead homogenizer (Bead Ruptor Elite v1.1, Omni International). Wounds were homogenized using two 15 sec cycles (with a 30 sec delay) at 7.1 ms^-1^, 4 °C. The tissue and beads were transferred into a glass extraction vial containing 1.25 ml hexane/isopropanol/glacial acetic acid (30:20:2). Tubes were rinsed with an additional 0.3 ml of buffer, then were vortexed and this was then added to the extraction vials. Internal standards (5 ng each of PC 14:0_14:0 and PE 14:0_14:0) and 5 ul of eicosanoid internal standards was added to each sample to give final concentration 75 nM (Supplementary Table 3). Following vortexing (1 min), hexane (1.25 ml) was added followed by vortexing, then centrifugation for 5 min at 1500 rpm, 4 °C. The upper organic phase was recovered into a clean tube. Hexane (1.25 ml) was added to the lower phase, which was again vortexed and centrifuged as above. The upper layer was combined with the previous hexane extract. A mixture of chloroform:methanol (1.9 ml, ratio 1:2) was added to the remaining aqueous phase. Following vortex, chloroform (0.625 ml) was added and samples again vortexed. HPLC grade water (0.625 ml) was added, and samples vortexed. Last, samples were centrifuged as above. The lower phase was carefully harvested and added to the hexane extracts obtained above. Samples were dried using a RapidVap (RapidVap, Labconco®) then resolubilized in methanol (200 µl), with careful vortexing. The samples were split in half (2 x 100 µl samples, sample A and sample B), with half being stored for eoxPL analysis, and the remainder analyzed for oxylipins (see below). Samples were stored at -80 °C until LC/MS/MS as described later.

#### Oxylipin extraction

100 µl methanol was added to sample B (listed above) followed by 1.655 ml HPLC-grade water. Glacial acetic acid (45 µl) was added to acidify and samples were mixed gently. Sep-Pak (C18 Waters) columns were loaded into a positive pressure (N_2_) manifold, then pre-conditioned using 100 % MeOH (12 ml), followed by acidified water (6 ml with 0.4 % glacial acetic acid). Samples were loaded onto the preconditioned columns and allowed to drip through using gravity. Acidified water (10 ml) was passed through the column followed by 6 ml hexane (under nitrogen pressure). Columns were allowed to dry for 30 min. Oxylipins were eluted from the column using methyl formate (8 ml) into glass extraction tubes then solvent was evaporated under vacuum. Lipids were reconstituted using methanol (100 μl) and stored at -80 °C until LC/MS/MS as described below.

#### LC/MS/MS analysis of oxylipins

Lipids were quantified using reverse phase LC/MS/MS. They were separated using a gradient of 30 – 100 % B over 20 min (A: water:mobile phase B 95:5 + 0.1% acetic acid, B: acetonitrile:methanol, 80:15 + 0.1% acetic acid) on an Eclipse Plus C18 Column (Agilent), and analyzed on a Sciex QTRAP^®^ 6500(5). Source conditions: TEM 475 °C, IS -4500, GS1 60, GS2 60, CUR 35. Chromatographic peaks were integrated using Multiquant 3.0.2 software (Sciex). The criteria for LOQ was signal:noise of at least 5:1 and with at least 5-6 points across a peak. Lipids were quantified using a standard curve generated and run at the same time as the samples and multiple reaction monitoring (MRM) channels and all assay parameters are provided in Supplementary Table 3 and(5). Each oxylipin was expressed per mg of skin tissue. Example chromatograms are provided in Supplementary Figure 10.

#### LC/MS/MS (reverse phase) analysis of oxidized phospholipids

Lipid extracts were separated using reverse-phase HPLC on a Luna 3 µm C18 150 × 2 mm column (Phenomenex, Torrance, CA) with a gradient of 50 – 100 % B over 10 min followed by 30 min at 100 % B (A: methanol:acetonitrile:water, 60:20:20 with 1 mM ammonium acetate, B: methanol, 1 mM ammonium acetate) with a flow rate of 200 µl min^−1^. Lipids were analyzed in MRM mode on a 6500 Q-Trap (Sciex, Cheshire, United Kingdom), monitoring transitions from the precursor to product ion (dwell 75 ms) with TEM 500 °C, GS1 40, GS2 30, CUR 35, IS − 4500 V, DP − 50 V, EP − 10 V, CE − 38 V and CXP at − 11 V. The peak area was integrated and normalized to the internal standard. For quantification of HETE-PEs, standard curves were generated with PE 18:0a/5-HETE, PE 18:0a/8-HETE, PE 18:0a/11-HETE, PE 18:0a/12-HETE and PE 18:0a/15-HETE synthesized as described previously(6). Information on MRM transitions and *m/z* values are presented in Supplementary Table 5. HETE-PEs were quantified using standard curves with DMPE used as internal standard, with LOQ at signal:noise 5:1. Due to the limited standards available, identifications for some lipids are putative, based on the presence of characteristic precursor and product ions, and retention times. Example chromatograms are provided in Supplementary Figure 11.

#### Chiral analysis of oxylipins

Lipid extracts were separated using a Chiralpak IA-U column (50×3.0 mm, Diacel) in reverse phase mode, with flow rate 300 ml/min, at 40 °C, according to(7), on a 6500 Q-Trap (Sciex, Cheshire, United Kingdom). Mobile phase A was water:0.1 % acetic acid, and B was acetonitrile:0.1% acetic acid, and the gradient was 10 % B raised to 100 % B over 20 min followed by a 2 min hold then decrease to starting conditions over 2 min. MRM transitions and instrument parameters were as used for oxylipin reverse phase analysis.

#### Gel zymography for MMP activity

On day 7, wounds were harvested, snap frozen in liquid N_2_ and stored at -80 ^O^C until processing. Tissue was homogenized using ceramic beads in a Bead Ruptor Elite v1.1 (Omni International) using two rounds at 8 m/sec for 15 sec with a 30 sec dwell time, in 0.3 ml ice cold lysis buffer with protease inhibitors (50 mm Tris-HCl, 150 mm NaCl, 1 % Nonidet P-40, 0.1 % SDS, 0.1 % deoxycholic acid, 2 μg/ml leupeptin, 2 μg/ml aprotinin, 1 mm PMSF pH 7.4). Samples were placed on a rotary carousel for 30 min at 4 °C. The homogenate was centrifuged at 15,000 × g for 5 min at 4°C. Protein was measured using the Bradford assay (Thermo fisher) and samples diluted to 15 μg/sample. Samples were diluted in sample buffer (Zymogram Sample Buffer, #1610764, Biorad) and loaded into the wells of precast gels (Novex™ 10% Zymogram Plus (Gelatin)) gels (Thermo Fisher)). Electrophoresis was performed with a Tris-glycine running buffer (LC-26754, Invitrogen), at 125 V for 140 min. The gel was incubated for 1 hr at room temperature in 3 % Triton X-100 on a rotary shaker, then incubated with development buffer (50 mm Tris base, 40 mm HCl, 200 mm NaCl, 5 mm CaCl_2_, and 0.2 % Brij 35) at 37 °C for 18 – 30 hr (depending on experiment) on a rotary shaker. Gels were stained using 0.5 % w/v Coomassie blue G-250 in 50 % DDH_2_0, 40 % methanol and 10 % acetic acid for 2 hr, and then destained for 1 hr using diluent (50 % DDH_2_0, 30 % methanol, 10 % acetic acid). Gelatinolytic activity was observed as clear zones or bands at the appropriate molecular weights. Mouse MMP-9 and human MMP-2 (R&D Systems) were used to locate bands. Bands were quantified using the gel analysis plugin on ImageJ.

#### Cell transfection and reporter assays

HEK293 cells were cultured and transfected as described previously with plasmids expressing mouse PPARγ and the *Firefly* luciferase under the control of 3x Ppar Responsive Element (PPRE)(8). The *Renilla* luciferase plasmid pRL-TK (Promega) was also included in the transfection as an internal control. At day 2, the cells underwent 24 h incubation in 50 μl media, with 1 μM rosiglitazone or DMSO (vehicle), lipid mixtures or methanol (vehicle). For each experiment (day) there were 4 replicates per condition and the experiment was repeated 3 independent times. On day 3, lysates were prepared, and a luciferase assay was performed using a Dual-Luciferase Reporter Assay System (Promega).

#### Statistical Analysis

To compare wounds using immunohistochemistry, Students T-test was used * p<0.05, ** p<0.01, *** p<0.001 and **** p<0.0001. For multi-time point analysis, data were analyzed using one-way ANOVA, with p < 0.05 considered statistically significant. For RNASeq, paired-end reads from Illumina sequencing were trimmed with Trimmomatic and assessed for quality using FastQC with default parameters. Reads were mapped to the Mouse GRCm38 reference genome using STAR and counts were assigned to transcripts using FeatureCounts with the GRCm38.84 Ensembl gene build GTF. Both the reference genome and GTF were downloaded from the Ensembl FTP site(9–14). Differential gene expression analyses used the DESeq2 package (14) to produce an excel output listing adjusted p-value and log2 fold change between conditions. The data were then filtered so that only genes with adjusted p-value < 0.05 were taken forward for analysis (Benjamini-Hochberg adjustment). Downstream pathway analyses and gene annotation were performed in ingenuity IPA (Qiagen IPA). Cytoscape was used to cluster genes by expression (FPKM) over all samples and timepoints(15). Pearson correlation coefficients were calculated for all possible gene pairs, and only highly significant genes retained (|r| > 0.8). During analysis, one wild-type (baseline) RNASeq sample failed a quality control check (*Alox15* expression data indicated it was a knockout) and was removed from further analysis. For temporal analysis of gene expression changes, lists of genes were generated using MATLAB_R2022a. A heatmap was generated using Clustergram in MATLAB, where data was standardized for each gene, so that the mean is 0 and the standard deviation is 1, and hierarchical clustering for rows performed. Data were analyzed using Ingenuity Pathway Analysis. For oxylipidomics, a two-way ANOVA was used to calculate differences within groups, (p < 0.05 considered significant), with Bonferroni post hoc test (https://statisty.app/two-way-anova-calculator). Heatmaps for oxylipins and oxPL were generated using an R script that processes the raw dataset, into log10 values, based on an average of 5 biological replicates, which then passes the resulting data to pheatmap (https://cran.r-project.org/web/packages/Pheatmap) to produce the final image.

### Supplementary Results

#### Structural analysis of resolvinD5

While most SPM were not detected, a peak co-eluting with the resolvinD5 (RvD5, 7*S*,17*S*-diHDOHE) standard was seen (Supplementary Figure 12). RvD5 represents one stereoisomer of 4 possible 7,17-diHDOHEs. The lipid is described to originate from 12/15-LOX dependent formation of 17-HDOHE, followed by its further oxygenation by 5-LOX, following transcellular uptake of 17-HDOHE into leukocytes(16). However reverse phase LC/MS/MS is unable to fully separate these isomers, and it was not possible to obtain an MS/MS spectrum to compare with the standard, due to the low levels of the lipid present in wounds. To address this, secondary MRMs were next analyzed, with two arising from fragmentation at C7 (m/z 141, 199) and one at C17 (m/z 261, Supplementary Figure 12 A-D). For the synthetic RvD5, the three MRMs co-eluted at 10.06 min as expected (Supplementary Figure 12 C). In day 1 wound extracts, a lipid was detected at 10.06 min showing co-eluting ions for m/z 199 and 141, however the third MRM (m/z 261) eluted slightly earlier (Supplementary Figure 12 B,D). Both the putative RvD5, and other later eluting lipids that were detected using these MRMs were absent from *Alox15^-/-^* wound extracts (Supplementary Figure 12 E). Next, chiral analysis was undertaken, with synthetic RvD5 eluting at 7.23 min, with the expected MRMs also co-eluting (Supplementary Figure 12 F). Chiral analysis of the wound lipid extract showed several peaks eluting between 6.8 – 8 min (Supplementary Figure 12 G). Based on ion ratios, the large peak at 7.89 min is likely to be the same as the two seen around 11 min on reverse phase LC/MS/MS (Supplementary Figure 12 A,G). A very small peak at 7.22 min had the same retention time as RvD5 standard (Supplementary Figure 12 F,G) and similar ion ratios with the m/z 199 ion dominating. A recent study using a similar chiral separation method monitored RvD5 and its isomers after pre-isolating 7,17-diHDOHE, using the MRM m/z 359-141, and showed that RvD5 elutes slightly later than its isomers, 7*R*,17S, 7*R*,17*R* and 7*S*,17*R*-diHDOHEs(17). In mouse wounds earlier eluting peaks that could represent these isomers are seen. These have the same MRMs as RvD5, in particular the peak at 6.84 min (Supplementary Figure 12 G, inset) indicating oxygenation at C7 and C17. Several other lipids eluted just after the putative RvD5 (7.3-7.6 min), and these may represent additional related structures such as positional isomers (Supplementary Figure 12 G). Spiking the wound lipid extract with synthetic standard showed co-elution of RvD5 with the peak at 7.2 min (Supplementary Figure 12 H,I). Overall, the data suggest the wound may contain low levels of RvD5, together with other related isomers of 7,17-diHDOHE. As for reverse phase analysis, the putative RvD5 peak along with all the other lipids detected using chiral analysis were absent in *Alox15^-/-^* wounds indicating their dependence on the enzyme (Supplementary Figure 12 J). However, 7,17-diHDOHE (coeluting with RvD5) was present at ∼0.5 % the levels of 17-HDOHE. Considering this, and the presence of isomers, it is possible RvD5 may have originated from non-enzymatic secondary oxidation of 12/15-LOX-derived 17S-HDOHE to form 7*R*,17*S*-diHDOHE and 7*S*,17*S*-diHDOHE(RvD5) eluting at 6.84 and 7.22 min, respectively. Further studies are needed to establish the origin of the lipid in wounds, for example pharmacological/genetic inhibition of 5-LOX, and MS/MS analysis of the 7*R*,17*S*-diHDOHE epimer under our chromatographic conditions.

#### Comparison of temporal changes in gene expression suggest additional transcription activators regulated by Alox15 beyond PPARg include elf4, Cebpb and Tcf3

To further interrogate *Alox15^-/-^* wounds for transcriptional regulators beyond PPARγ, a temporal analysis was performed on the RNASeq data. Here, analysis of individual strains separately allowed testing for genes behaving differently during progression of wound healing. In WT mice, 1705 transcripts significantly increased > 50 % on Days 0 and 4, while by Day 7, they reduced by > 25 % compared to Day 4 (Supplementary Figure 13 A, List 1). Thus, these elevate on acute injury, then return close to normal after one week (Supplementary Table 9). Interrogating these in *Alox15^-/-^* mice, 154 did not increase on Day 4 by > 25 % compared to Day 0 (Supplementary Figure 13 B, Supplementary Table 10). Thus, these failed to elevate during acute inflammation in *Alox15^-/-^*. Using IPA analysis, several transcription factors were identified as possible upstream regulators. *Elf4* is a known anti-inflammatory transcription regulator of inflammation, which targets several genes in the list, including *Anln, Asf1b, Ccnb2, Cdca3, Cenpa, Cenpe, Cks2, E2f8, Hmmr, Kif4a, Mcm10, Ndc80, Oip5, Rrm2, Tpx2*(*18*). In support of this idea, we found that expression of *Elf4* was significantly increased by wounding in both WT and *Alox15^-/-^* mice (Supplementary Figure 13 D). This indicates that while *Elf4* is upregulated by wounding, it may not be transcriptionally active in the absence of 12/15-LOX. Additional transcription regulators strongly associated with the initial response to wounding included *Tcf3* and *Cebpb*. *Tcf3* promotes cell migration and wound repair (19), and *Cebpb* is involved in macrophage repair responses and inflammation (20, 21). Expression data for these genes showed *Cebpb* is upregulated on Day 4, significantly in *Alox15^-/-^*, while reduced back to baseline expression by Day 7. However, *Tcf3* expression was unaffected by wounding in either strain (Supplementary Figure 14 A,B).

Last, genes in List 1 were re-interrogated to identify transcripts that reduced < 25 % on Day 7 compared to Day 4 in *Alox15^-/-^*. These represent genes that fail to resolve to basal levels during inflammation in *Alox15^-/-^*. Here, 538 were identified (Supplementary Figure 14 C, Supplementary Table 11). IPA analysis of these proposed “lipopolysaccharide” as the top upstream regulator, consistent with the failure of many known pro-inflammatory genes which respond to this bacterial product to reduce back to basal levels as shown in our earlier analysis, e.g., *Il6*, *Ptgs2*, and *Il1b* (Supplementary Figure 7). Similarly, IPA also proposed the top affected canonical pathway for List 3 as “Pathogen Induced Cytokine Storm Signaling” which includes 33 genes which failed to fully resolve. These are shown in a heatmap, comparing Days 4 or 7 with Day 0 in both WT and *Alox15^-/-^* mice. Three distinct groups are seen (Supplementary Figure 14 C):

i. Genes that elevate by Day 4 and then are reduced by Day 7 in WT. They elevate to a similar level in *Alox15^-/-^*but do not fall back to baseline by Day 7 or elevate further by that time (*Fos, Ddx58, Stat1, Ccr3, Nlrc4, Naip1, Faslg, Gsdmd, Clec7a, Itb, Lif, Stat4)*
ii. Genes that elevate by Day 4 then reduce by Day 7 in WT, while elevating higher in *Alox15^-/-^*at Day 4, and not falling back to baseline by Day 7 (*Cxcl3, Aim2, Ccl4, Ccl3l3, Cxcl10, Pgf, Nos2, Cxcl2, Tlr2, Mlkl, Sting1, Cxcr4, Csf2rb, Zbp1, Cklf*)
iii. Genes that elevate far higher in WT than *Alox15^-/-^* but reduce back by Day 7 in both (*Il1r1, Ccl7, Cxcl6, Col13a1, Ccr5, Irf7*).

Analysis using STRING 11.5 showed that group (i) genes are members of networks that regulate cytokines, including IFN-I, IL-12, IL-21, IL-35, IL-20 family and IL-6 family signaling networks. For groups (ii) and (iii) the main KEGG pathways were “cytosolic DNA-sensing” and “IL-17 signaling”, respectively. Notably this analysis confirms our earlier data which indicates that inflammatory signaling is strongly impacted by *Alox15*^-/-^ deletion, while identifying a large number of novel targets for further study.

### Supplementary Figure Legends

**Supplementary Figure 1.**
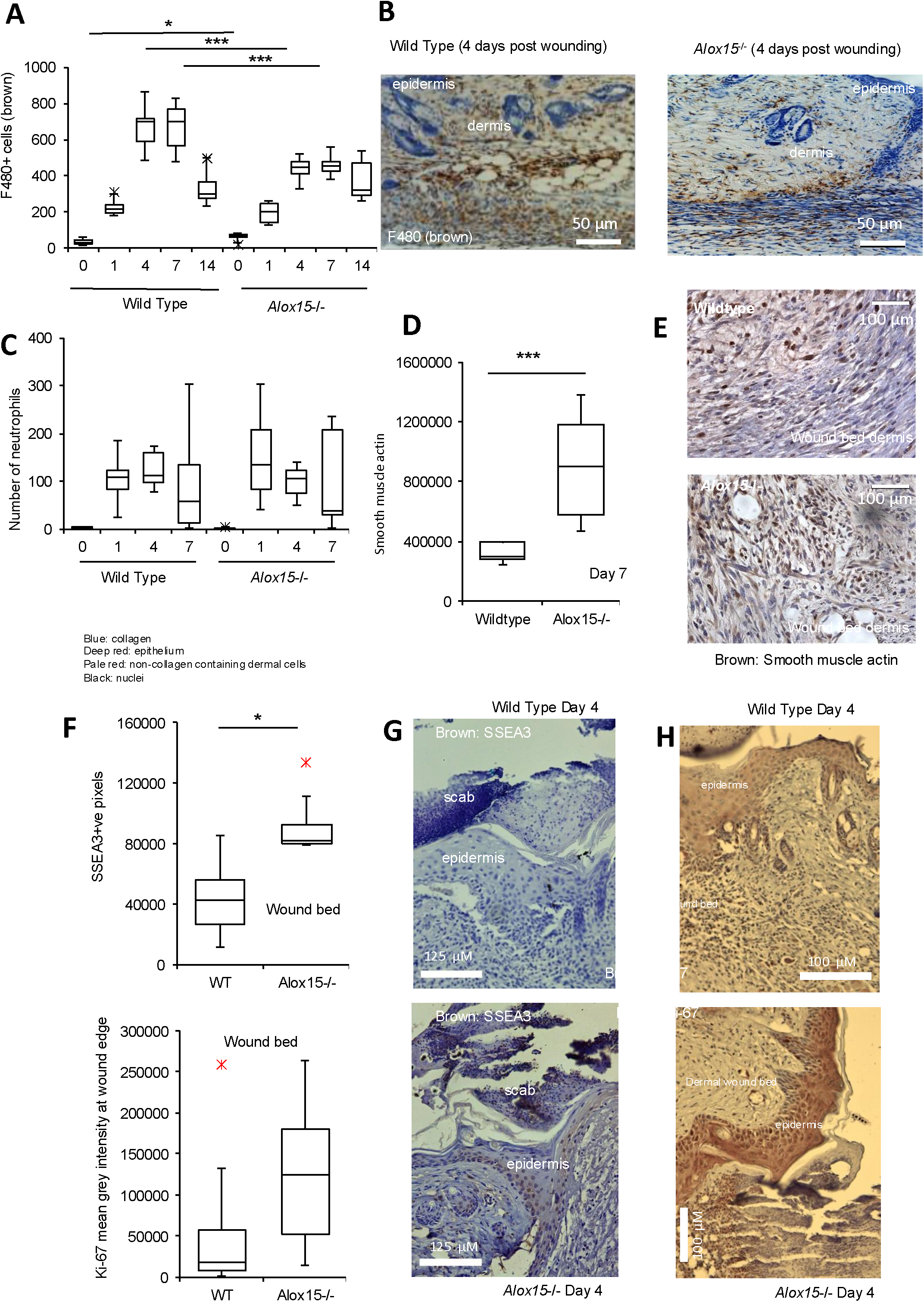
*Alox15*^-/-^ wounds show reduced wound bed macrophages, and increased smooth muscle actin, SSEA3 and Ki-67 expression. *Panel A. Alox15 deletion leads to reduced macrophage influx*. F480+ve cells were measured in wounds as described in Methods (n= 6–8/group, 4-9 fields per wound). *Panel B. Representative images from Panel A*. *Panel C. Neutrophil cell numbers (visualized with by Ly6g DAB positive staining) are similar in wildtype and Alox15^-/-^ mice*, (n = 7–8/group, 4-9 fields per wound). *Panel D. The myofibroblast marker smooth muscle actin alpha was increased in Alox15^-/-^ wounds*. Cells were stained with anti-alpha smooth muscle actin and visualized with DAB staining (n = 7- 8/group). For panels B,H: unpaired t-test, * p < 0.05, *** p<0.005. *Panel E. Representative images of smooth muscle actin. Panel F. SSEA3 and Ki-67 are elevated in SSEA3 and Ki-67 Alox15^-/-^ wounds*. Expression was measured using immunohistochemistry, followed by DAB visualization. n = 6/group (SSEA3), 10/group (Ki67). *Panels G,H. Representative data from Panel F*.

**Supplementary Figure 2.**
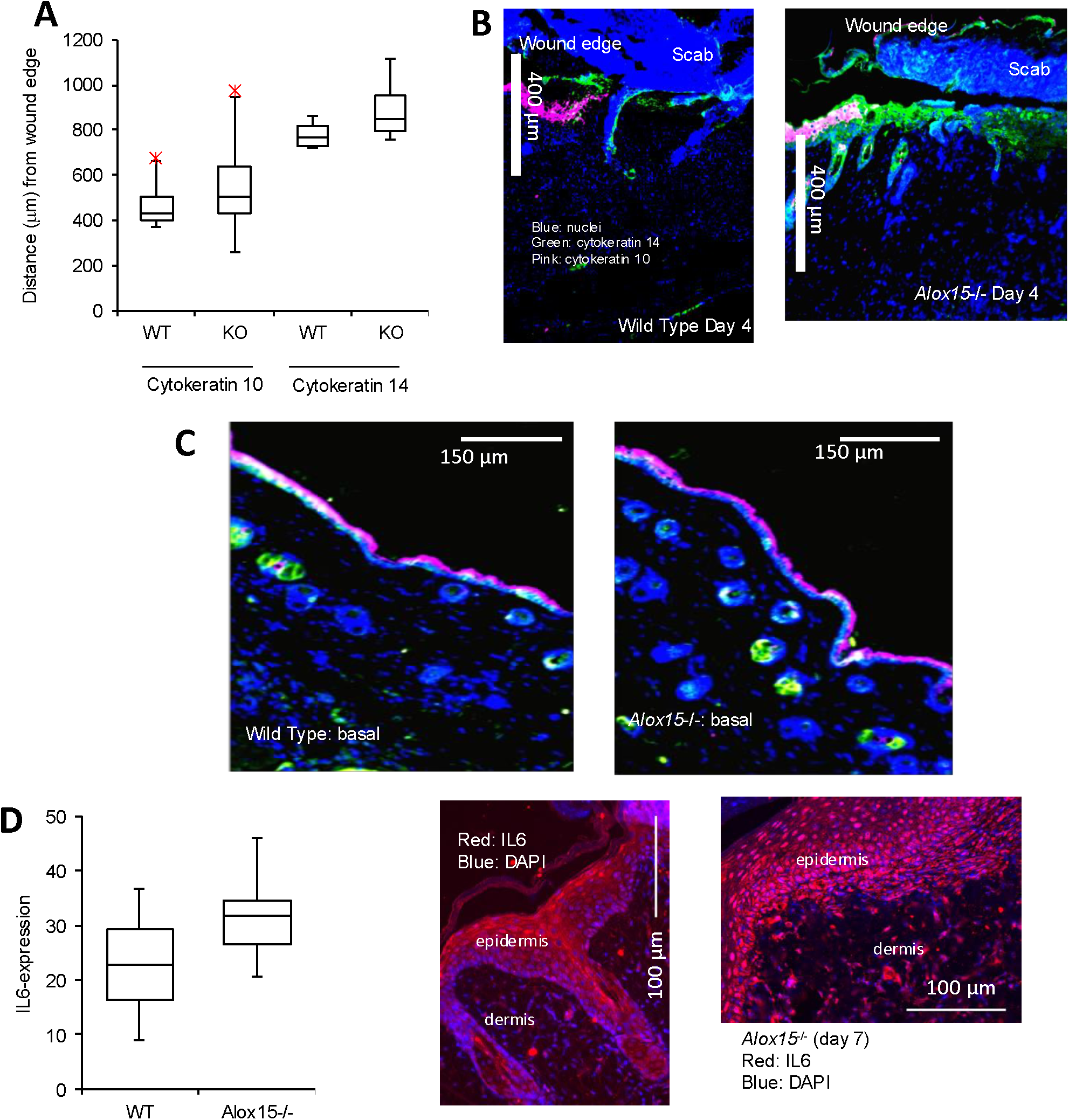
Epithelial proliferation and terminal differentiation of keratinocytes are not impacted in *Alox15^-/-^* either basally or post-wounding, and IL-6 is unaffected. *Panel A. Cytokeratin 14 migration migrated from the wound edge in Alox15^-/-^ wounds at day 4 is not impacted.* Quantification of the migratory distance of highly proliferative non-differentiated cytokeratin 14 (green) and non-proliferative terminally differentiated cytokeratin 10 (pink) was quantified using fluorescence immunohistochemistry. n = 4/group. *Panel E. Representative images from Panel B. Panel C. Representative images showing cytokeratin 10 and 14 staining in non-wounded skin*. All panels, unpaired Students t-test, * p<0.05, *** p, 0.005. *Panel D. Alox15^-/-^ wounds show unchanged IL-6 expression*. IL-6 was measured using fluorescence immunohistochemistry. n = 6/group.

**Supplementary Figure 3.**
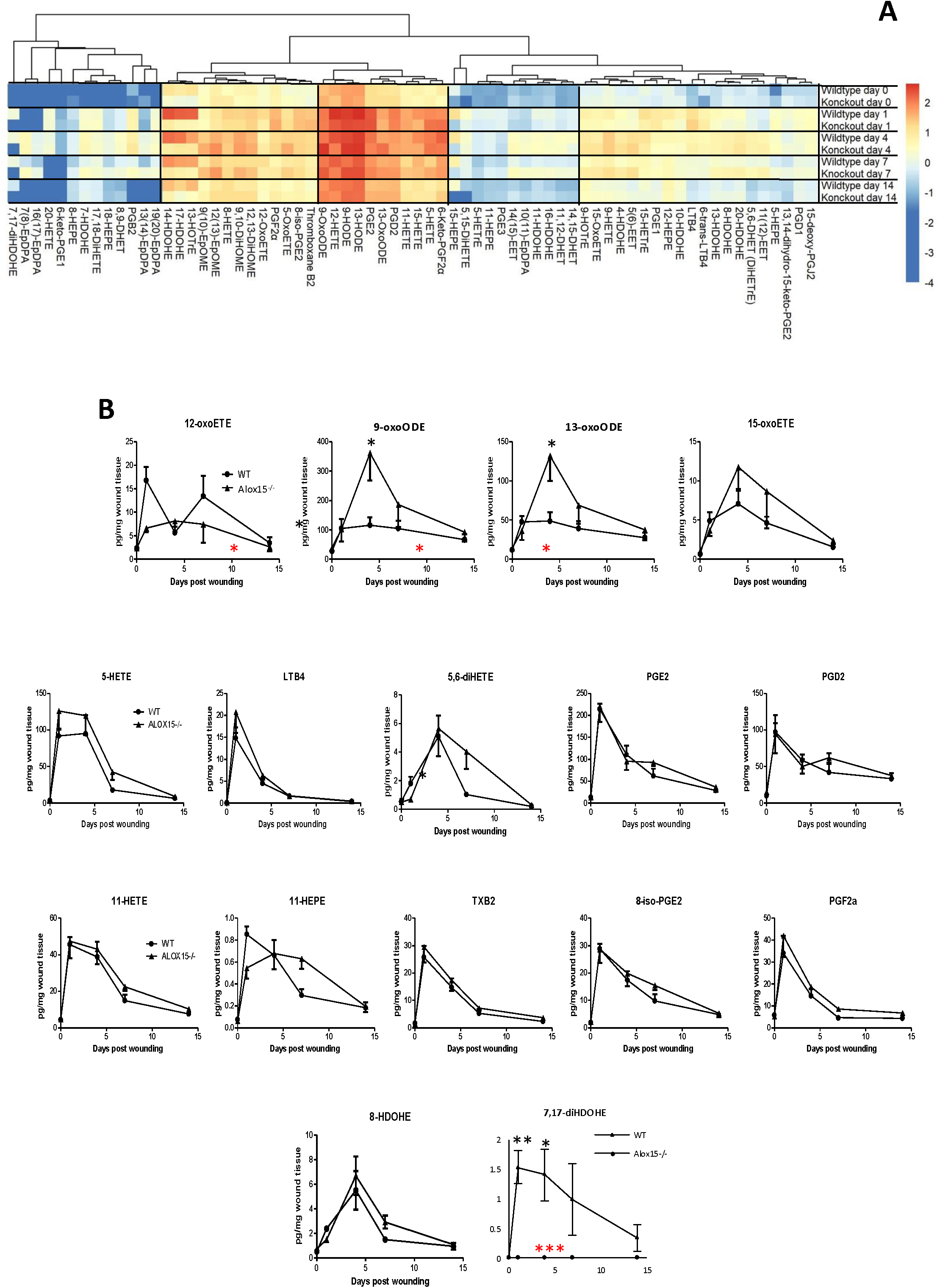
Heatmap and time course data for oxylipin levels during wounding shows that *Alox15^-/-^*wounds generate lower levels of many oxylipins. *Panel A*. *A heatmap shows log10 of mean values for all lipids across all groups tested.* Oxylipins were measured using LC/MS/MS as outlined in Methods (n = 5 samples/time point, with 4 wounds pooled/sample). *Panel. Oxylipins are rapidly elevated post-wounding, but many are reduced in Alox15^-/-^ wounds.* Oxylipins were measured using LC/MS/MS as outlined in Methods. n = 6 samples/time point, with 4 wounds pooled/sample. For all panels differences between groups were analyzed using two-way Anova (red stars), with Bonferroni post hoc test between individual time points (black stars), mean ± SEM, * p < 0.05, ** p<0.01, *** p < 0.005.

**Supplementary Figure 4.**
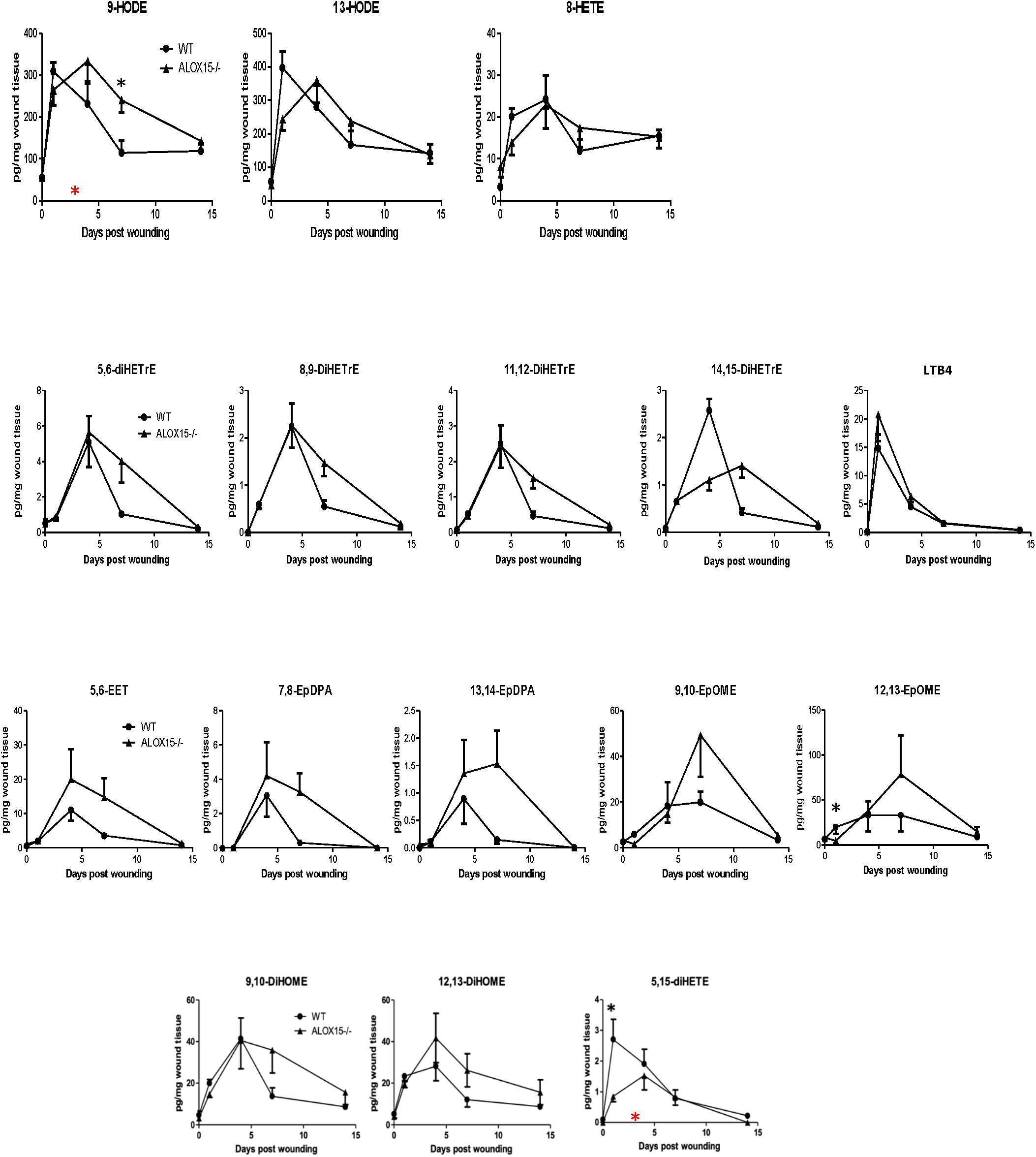
Time course data for oxylipins generated during wounding, as in Supplementary Figure 2. Oxylipins were measured using LC/MS/MS as outlined in Methods. n = 5 samples/time point, with 4 wounds pooled/sample. For all panels differences between groups were analyzed using two-way Anova (red stars), with Bonferroni post hoc test between individual time points (black stars), mean ± SEM, * p < 0.05, ** p<0.01, *** p < 0.005.

**Supplementary Figure 5.**
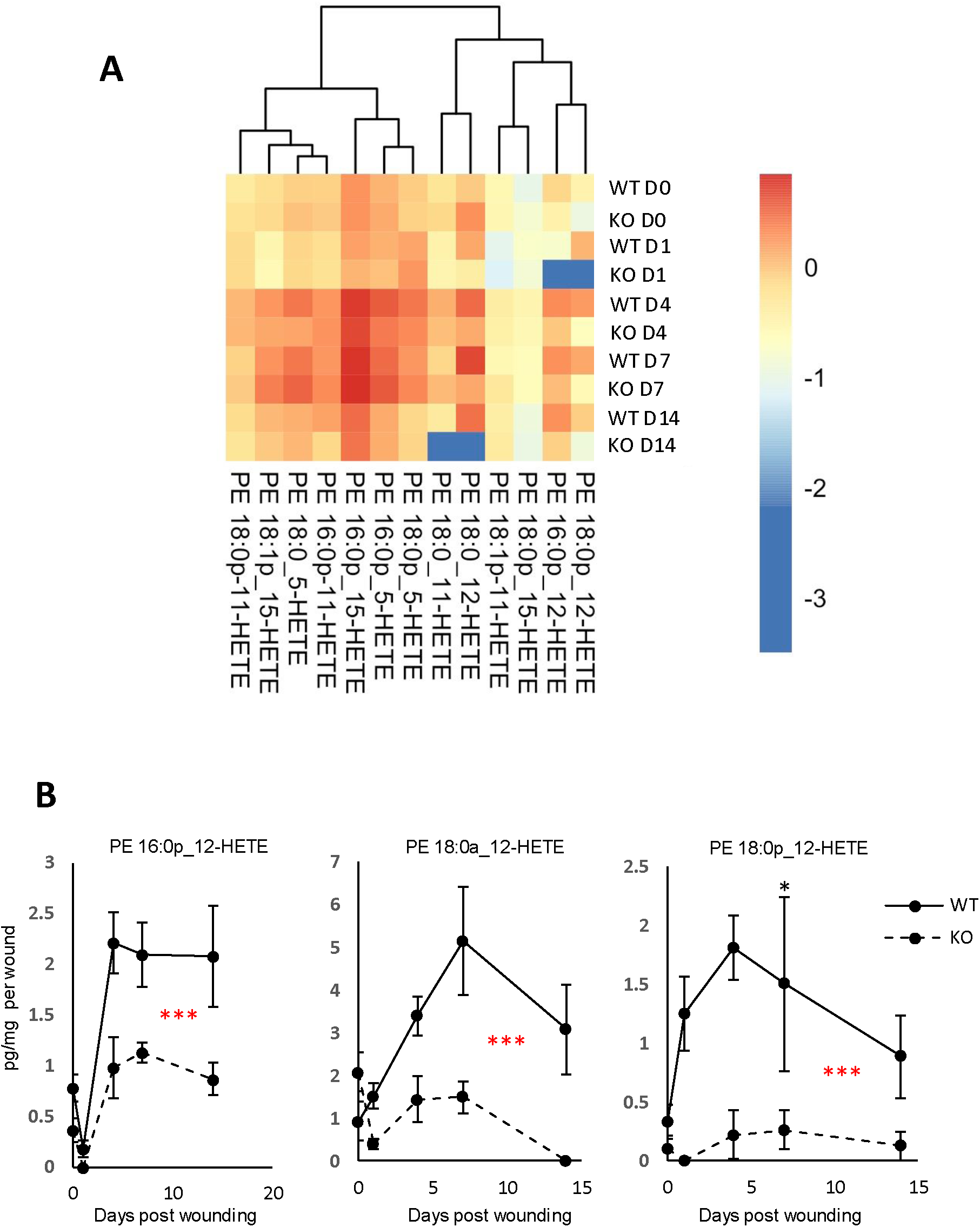
Heatmap of eoxPL generation during wounding, and individual timecourses of 12-HETE-PEs. *Panel A.* Heatmap shows expression (log10) of mean values for all lipids across all groups tested eoxPL were measured using LC/MS/MS as outlined in Methods. n = 5 samples/time point, with 4 wounds pooled/sample. *Panel B. 12-HETE-PE isomers are significantly reduced in Alox15^-/-^ wounds.* Oxidized phospholipids were measured using LC/MS/MS as outlined in Methods (n = 5 samples/time point, with 4 wounds pooled/sample). Unpaired t-test, * p <0.05, ** p < 0.01, *** p < 0.005.

**Supplementary Figure 6.**
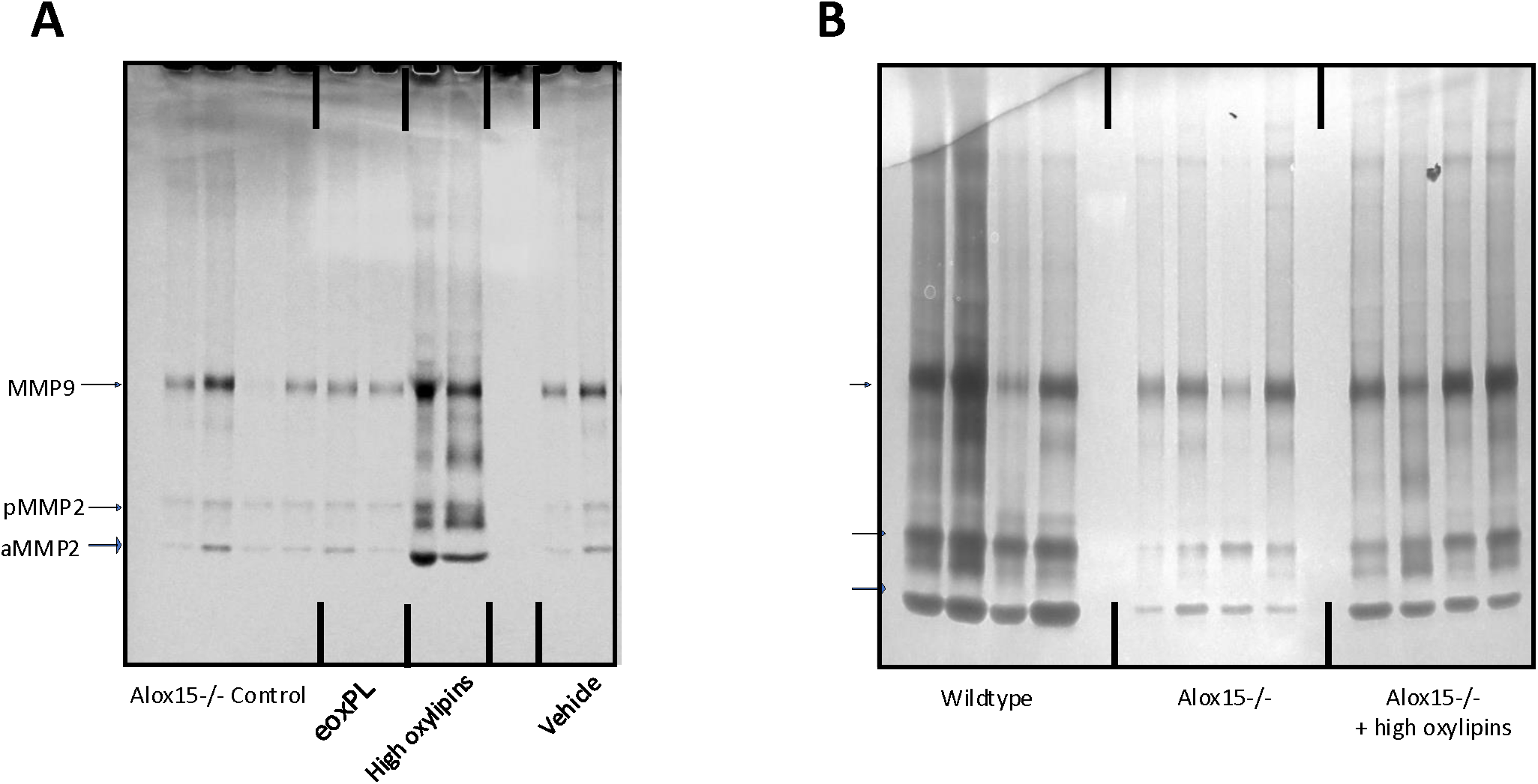
Raw data for MMP quantitation in wounds *Panels A,B.* Gel zymography showing data used to quantify MMP activities during wounding.

**Supplementary Figure 7.**
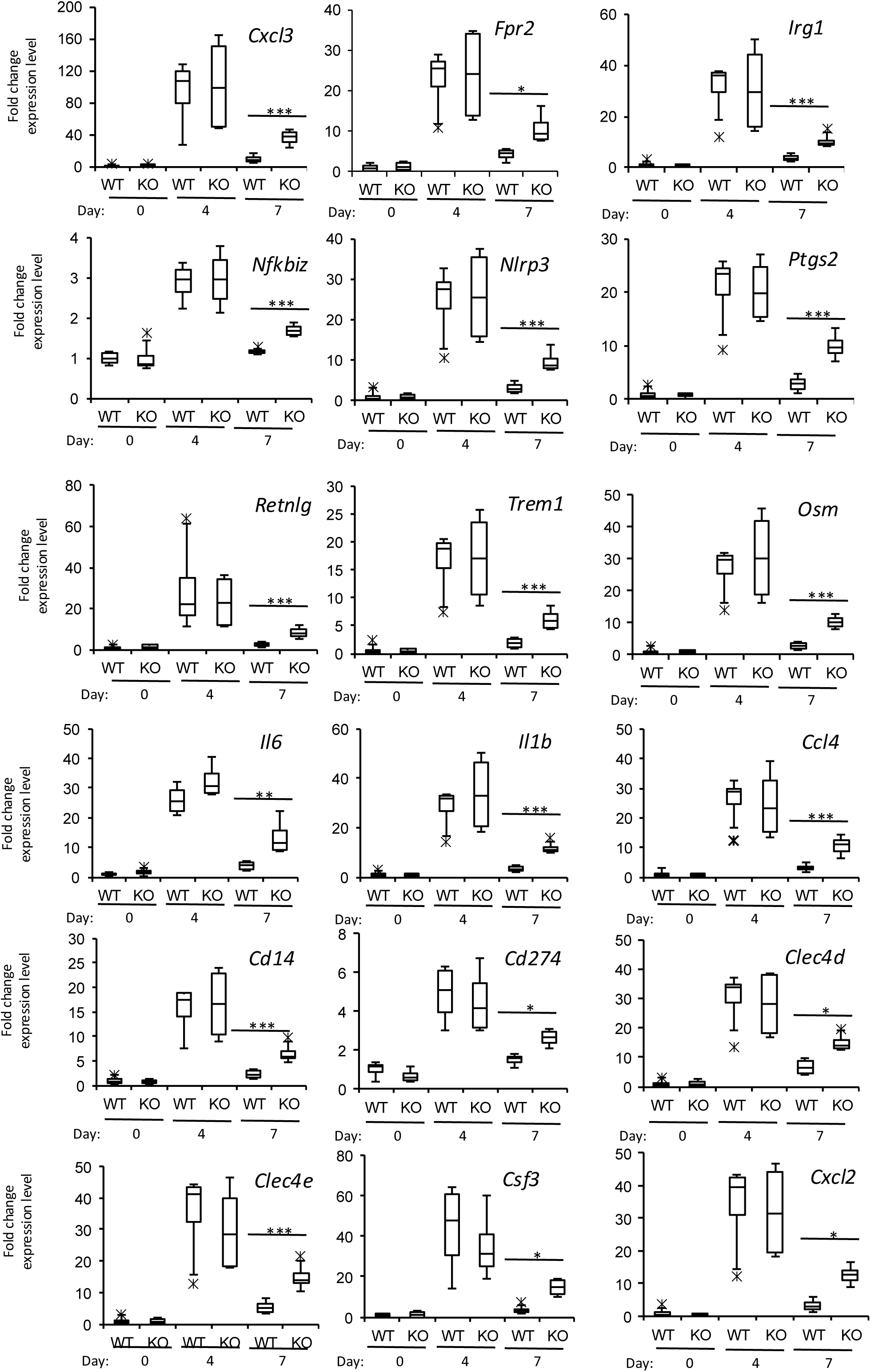
Genes in the network show the same pattern of expression throughout the time course, increasing on Day 4, but with a failure to revert to baseline at Day 7 in *Alox15^-/-^*wounds. Data for several affected genes are shown, all gene expression data was normalized to its Day 0 mean value, and then expressed as fold-change (n = 3 – 4 per group). For all gene expression data, students t-test, followed by Benjamin Hochberg correction: * p <0.05, ** p < 0.01, *** p < 0.005.

**Supplementary Figure 8.**
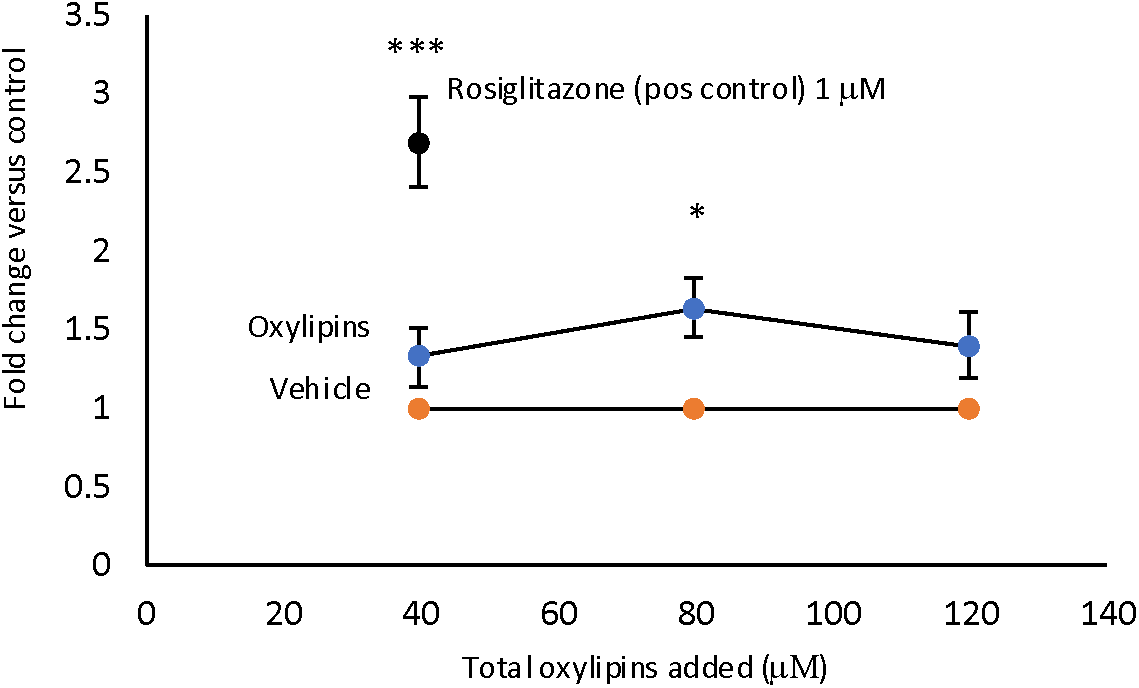
High oxylipins activate PPARγ transcription. Oxylipins, Rosiglitazone or vehicle controls were added to HEK293 cells expressing mouse PPRE, as described in Methods. After 24 hrs, luciferase activity was analyzed. Data are shown normalized to the relevant vehicle control (n = 3 independent experiments, mean +/- SEM, students t-test, * p <0.05, *** p < 0.005.

**Supplementary Figure 9.**
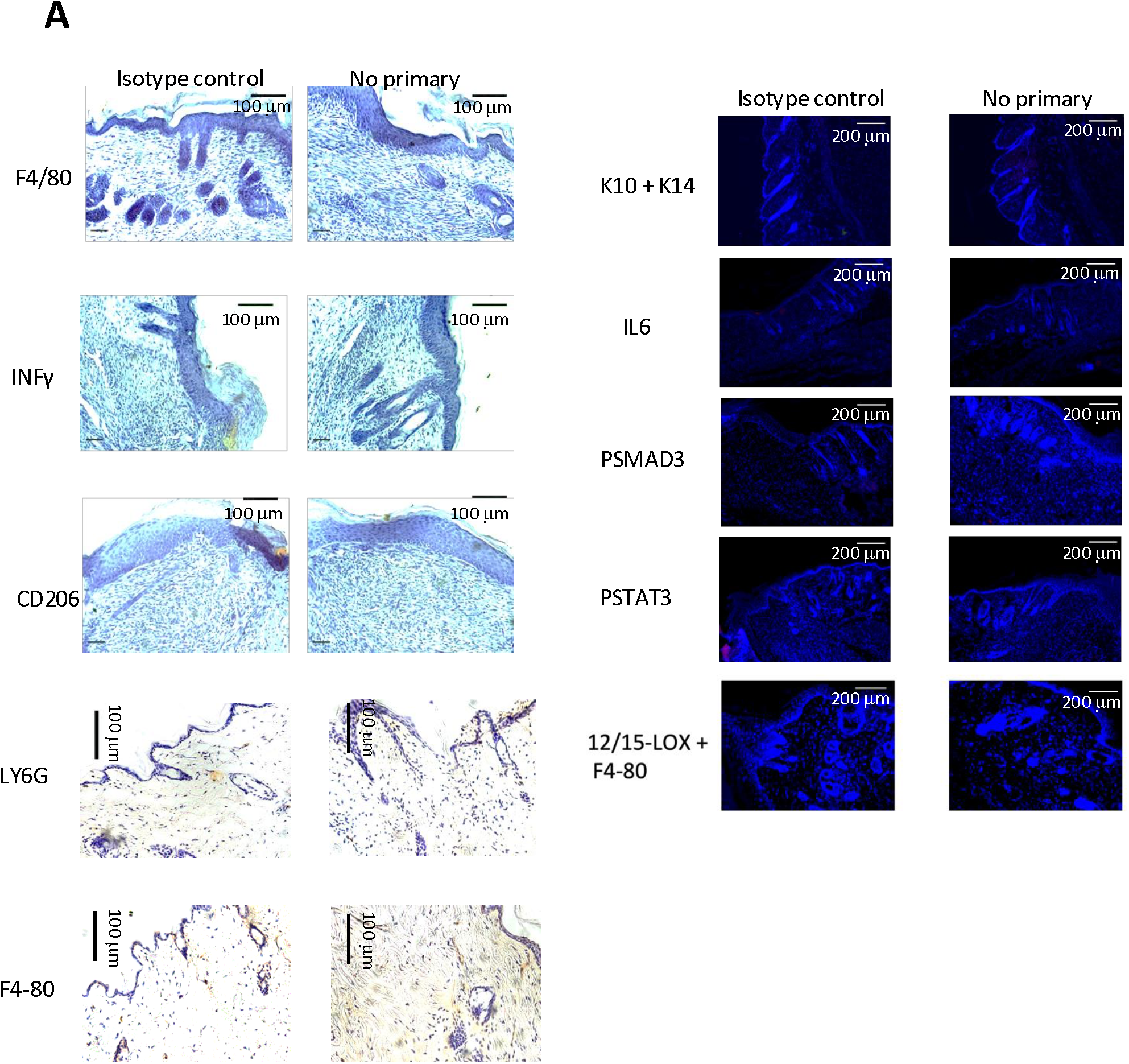
Isotype and no-primary controls for antibodies in the study. Left panels: To confirm target-specific DAB staining both a no primary control and isotype control were run for the antibodies F4/80, INFγ and CD206 followed by hematoxylin counter staining. Images were taken at 20x magnification, scale bar =100μm. Right panels: To confirm target specific immunofluorescence staining both a no-primary control and isotype control was run for the antibodies, Cytokeratin 10 (K10), cytokeratin 14 (K14), IL6, Phospho SMAD3, Phospho STAT3, followed by a DAPI stain. Images were taken at 10x magnification, scale bar =200μm.

**Supplementary Figure 10.**
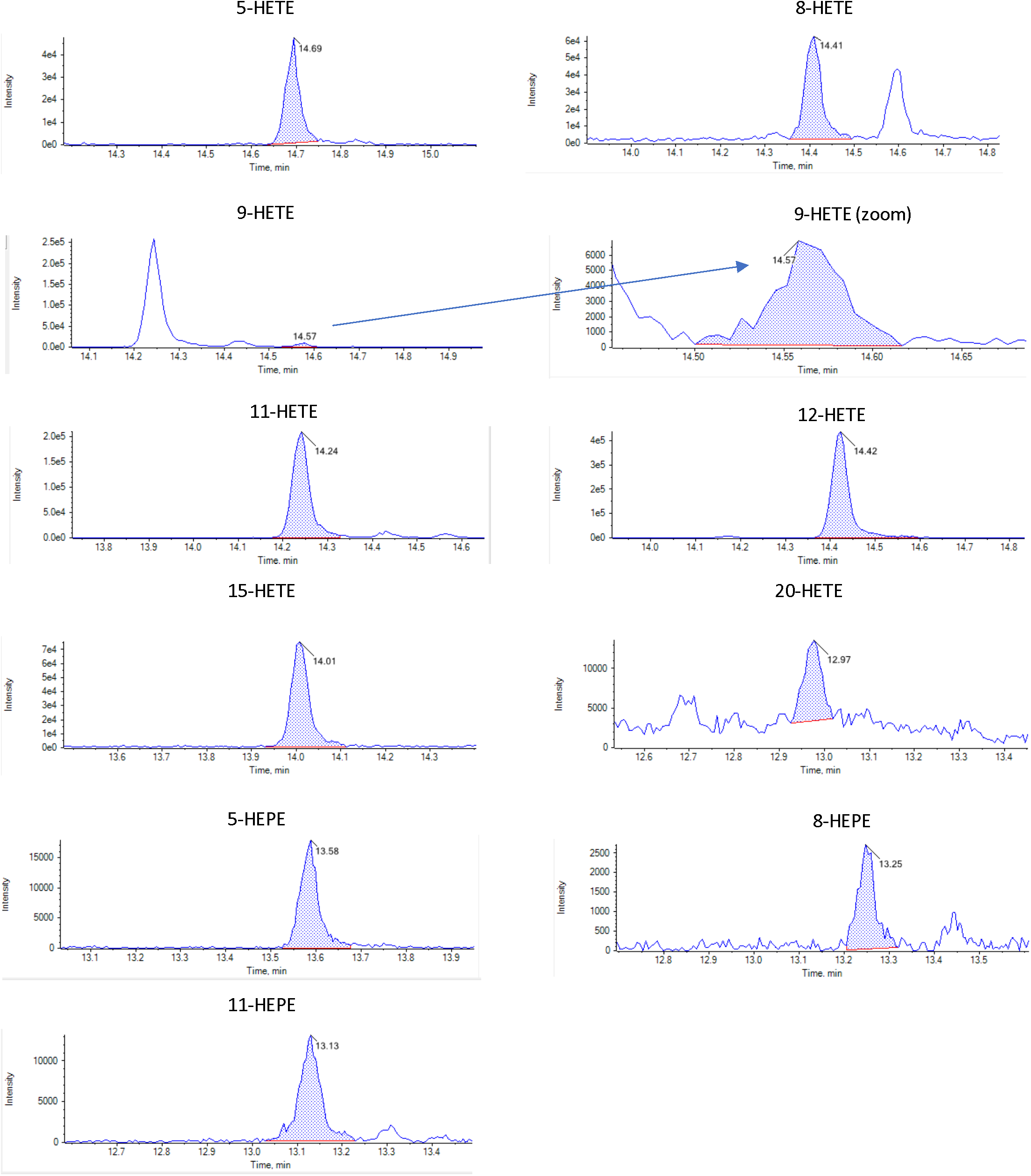
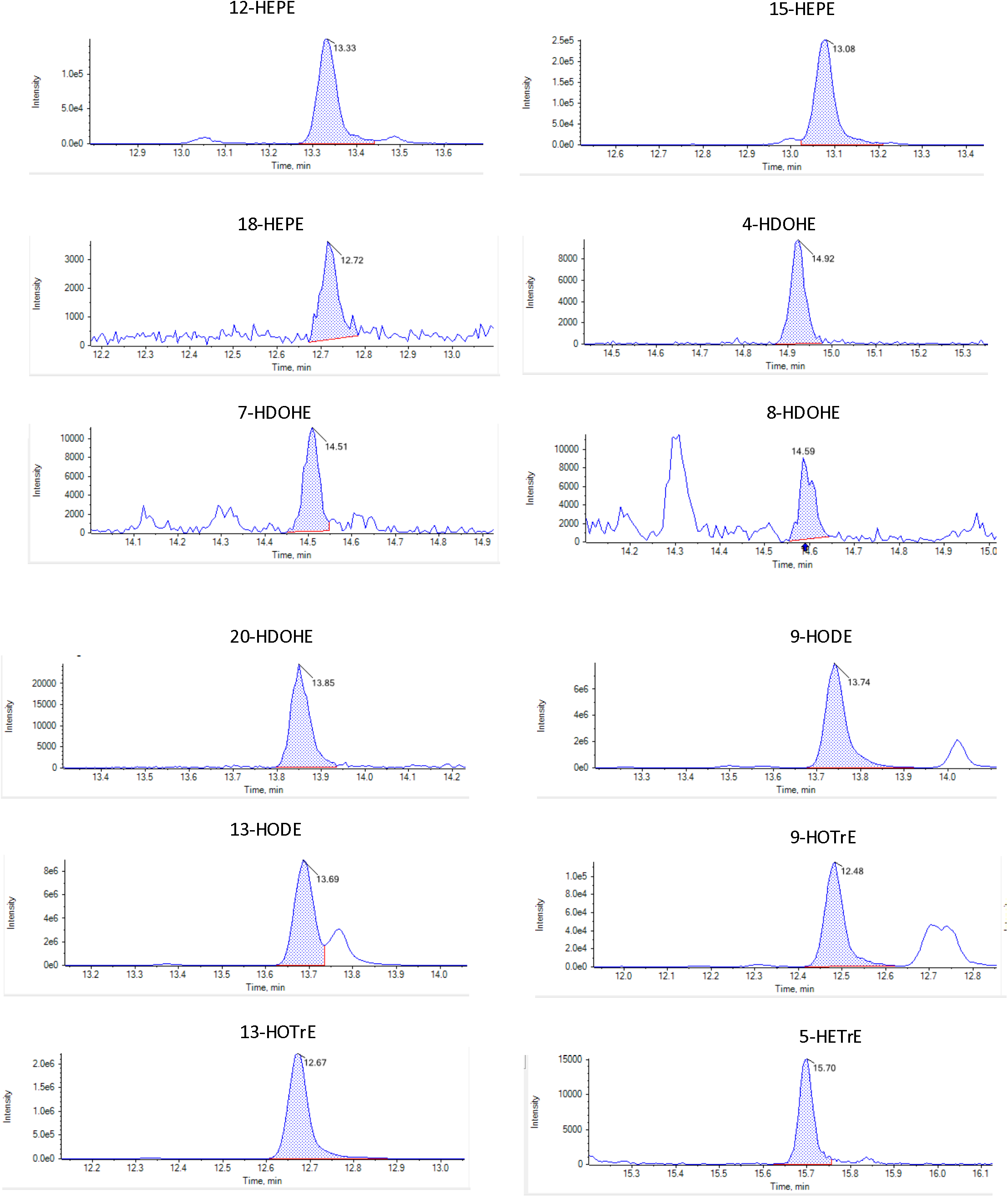
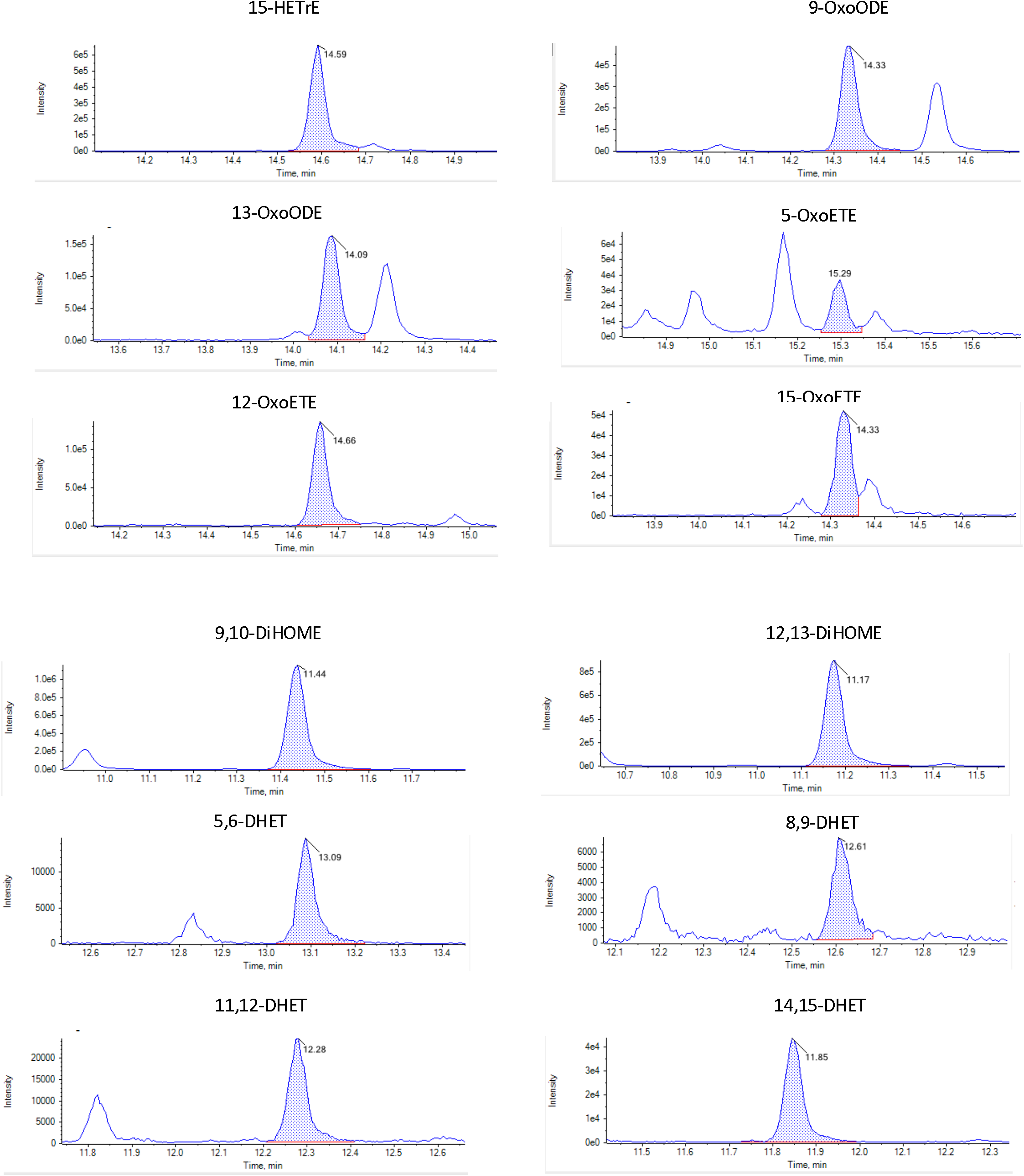
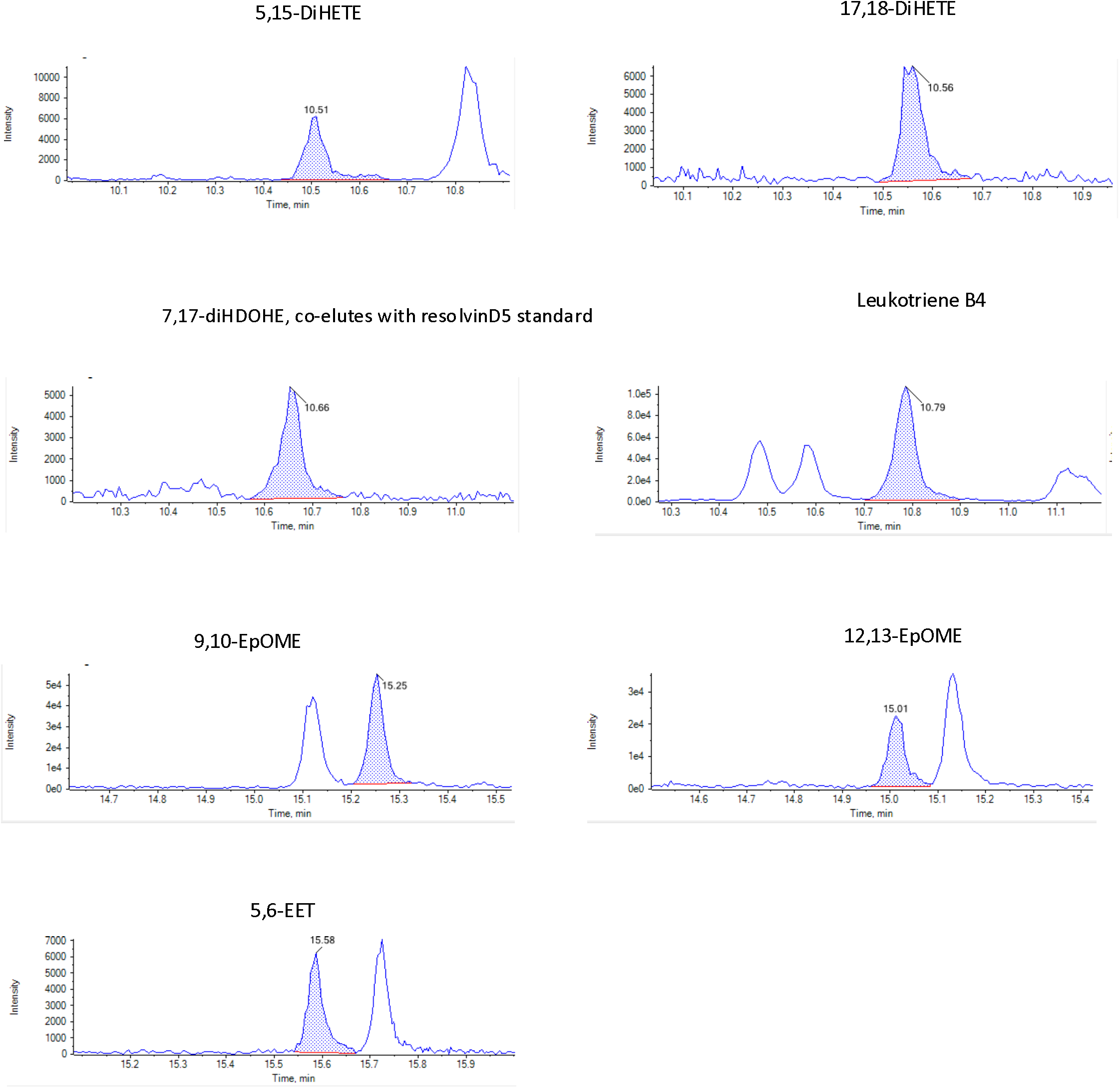
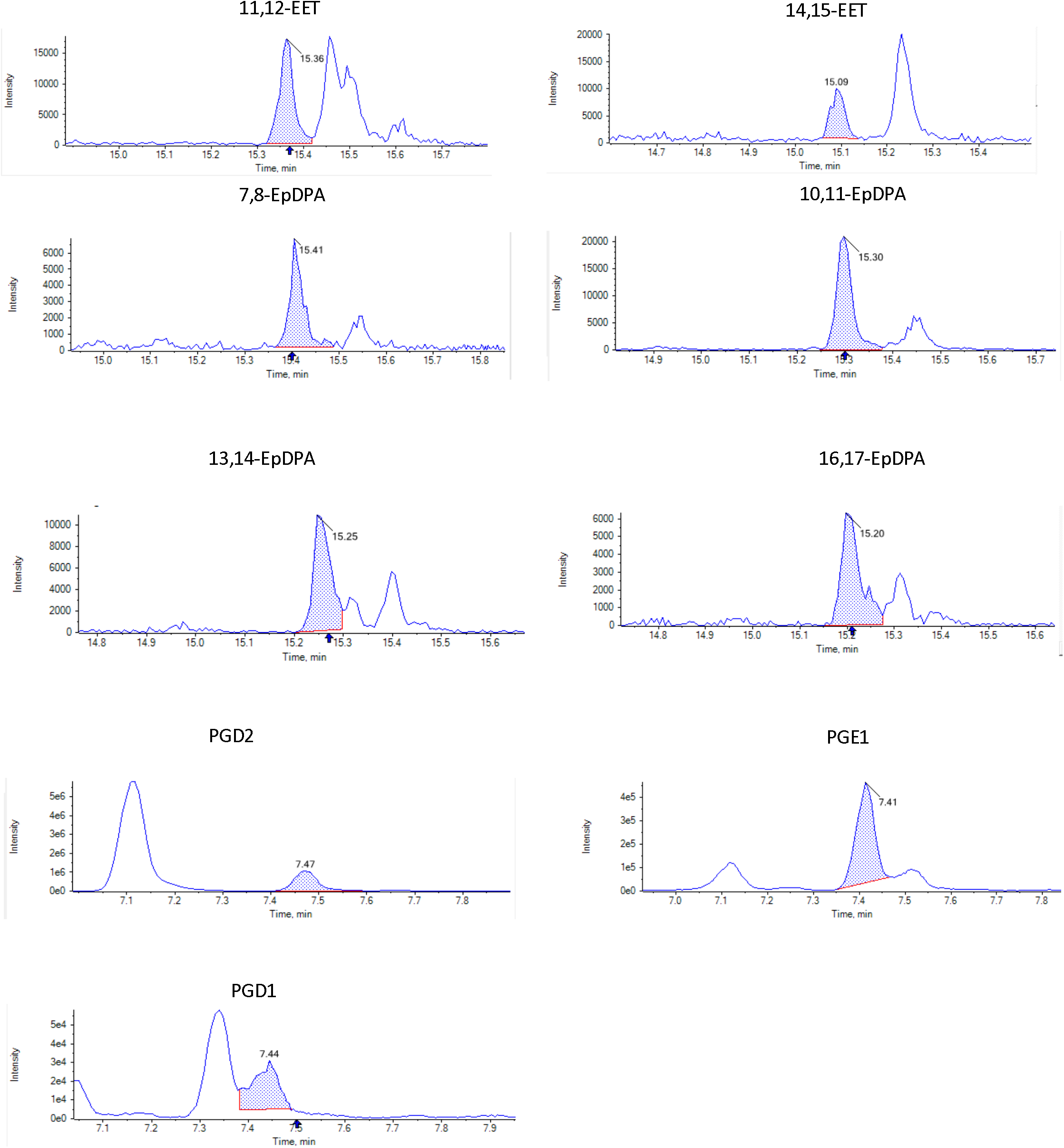
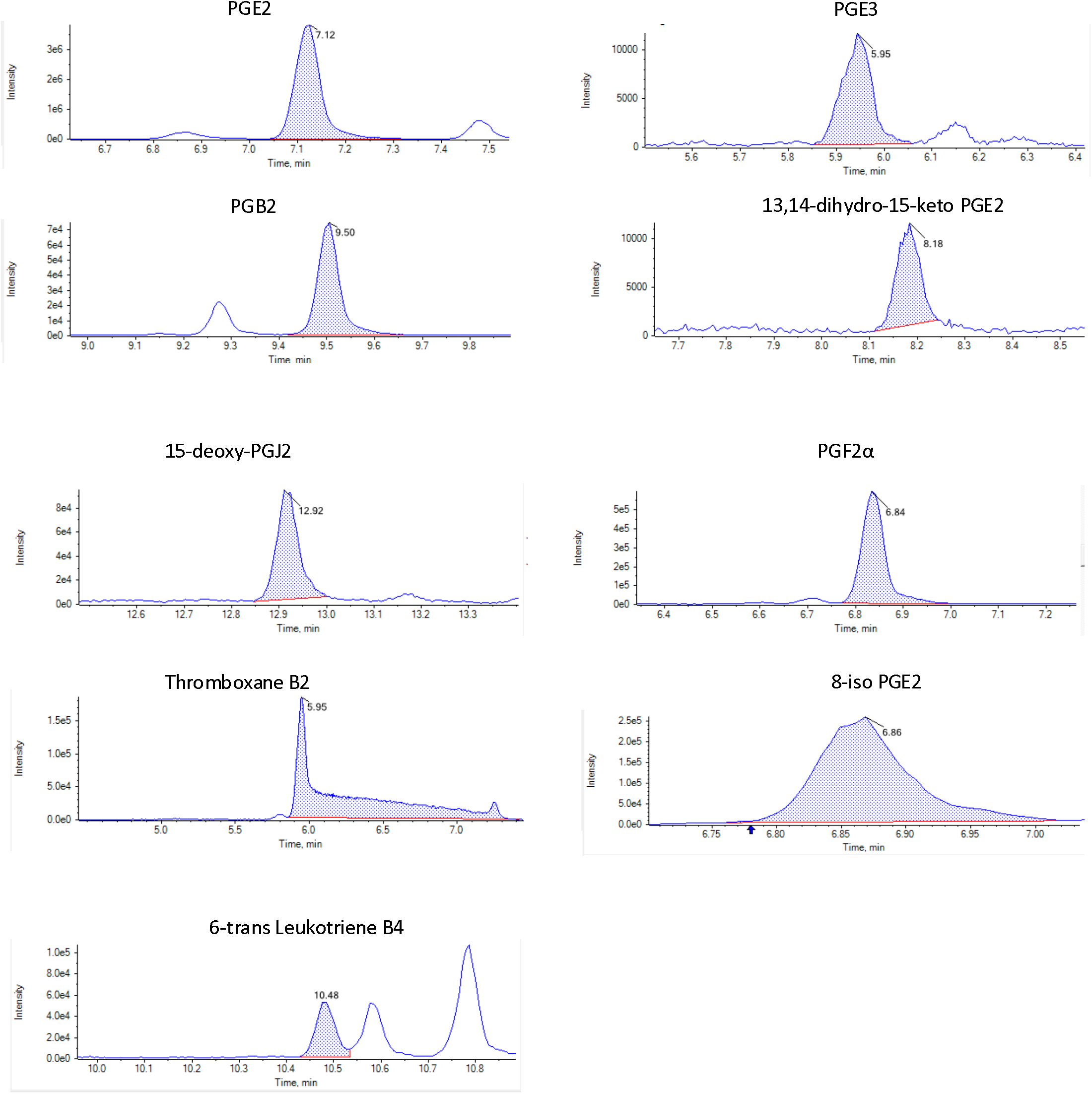
Representative chromatographic peaks for oxylipins detected during wounding. Screenshots were taken from Multiquant software, with the shaded area indicating the peak integrated. Wound lipids were confirmed to co-elute with primary standards in the same analytical batch.

**Supplementary Figure 11.**
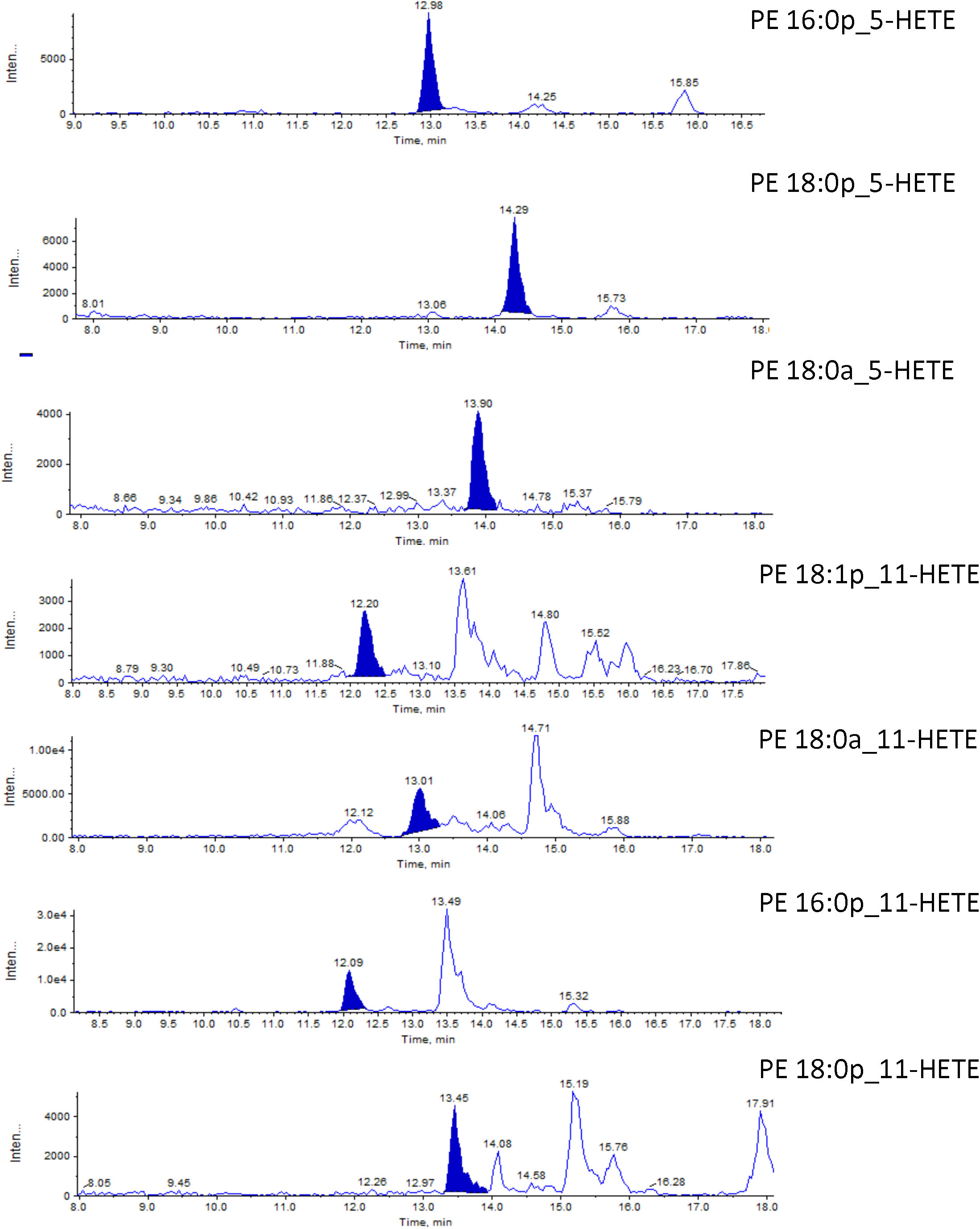
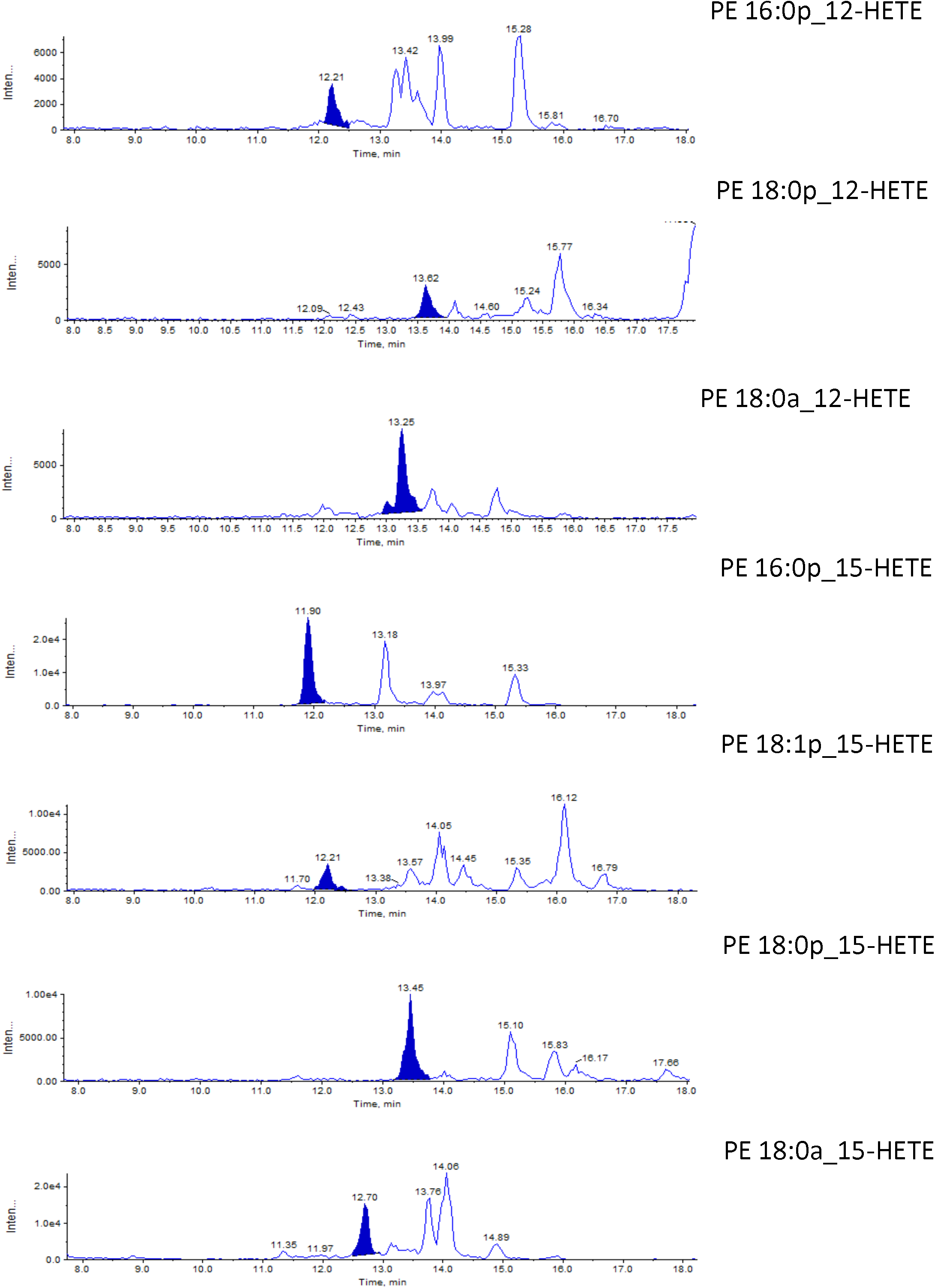
Representative chromatographic peaks for eoxPL generated during wounding. Identity was verified comparing retention time with standards as outlined in (6), based on comparison with PE 18:0a_HETE for the relevant positional isomers. Note that standards for 16:0p, 18:0p and 18:1 forms are not available, and so relative RT compared to standards is used along with MRM transitions which use internal daughter ions for all HETE positional isomers, along with LOQ of > 5 for signal:noise for peaks. The order of elution is characteristic for the different sn1 forms, as shown in (22–24).

**Supplementary Figure 12.**
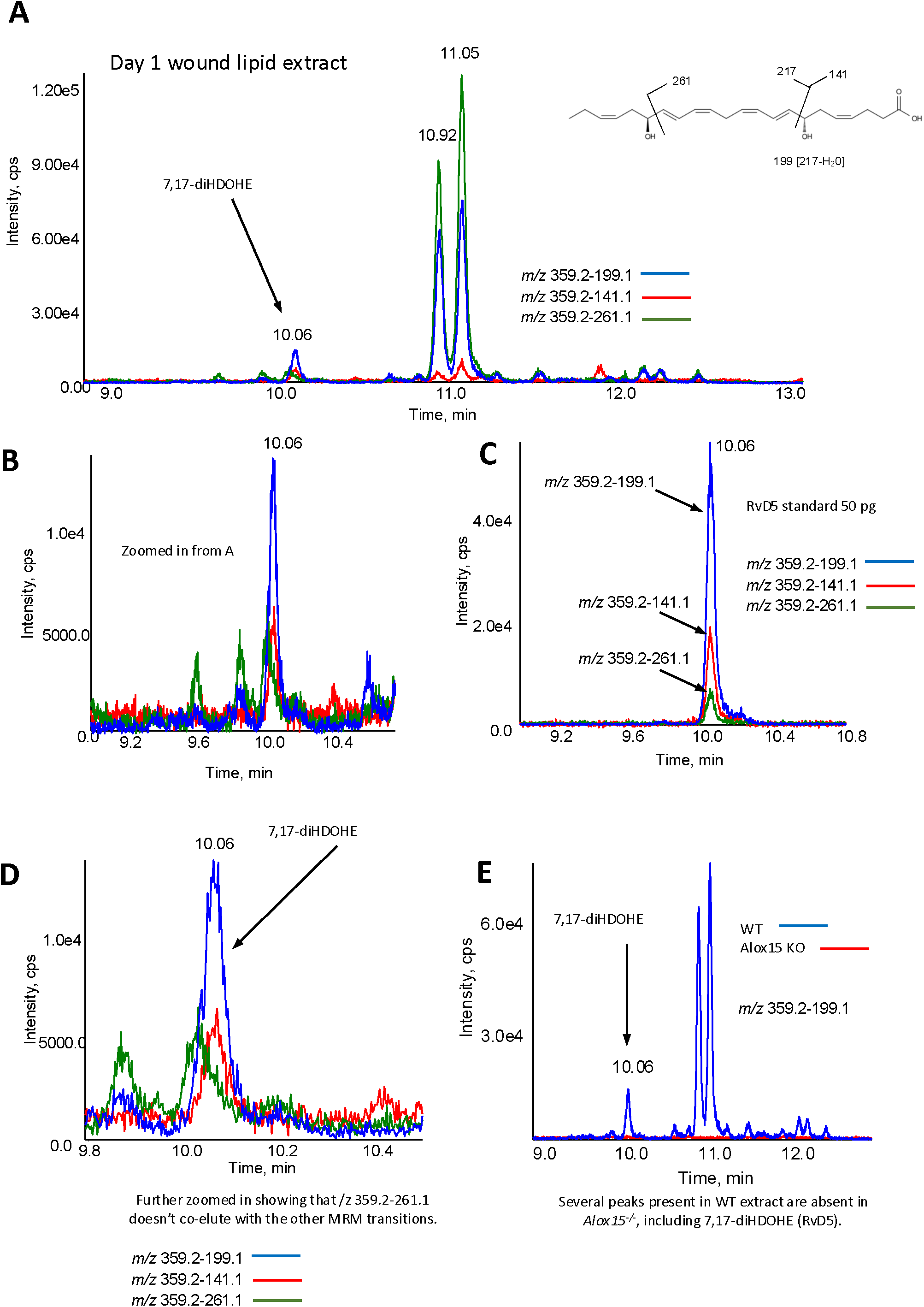
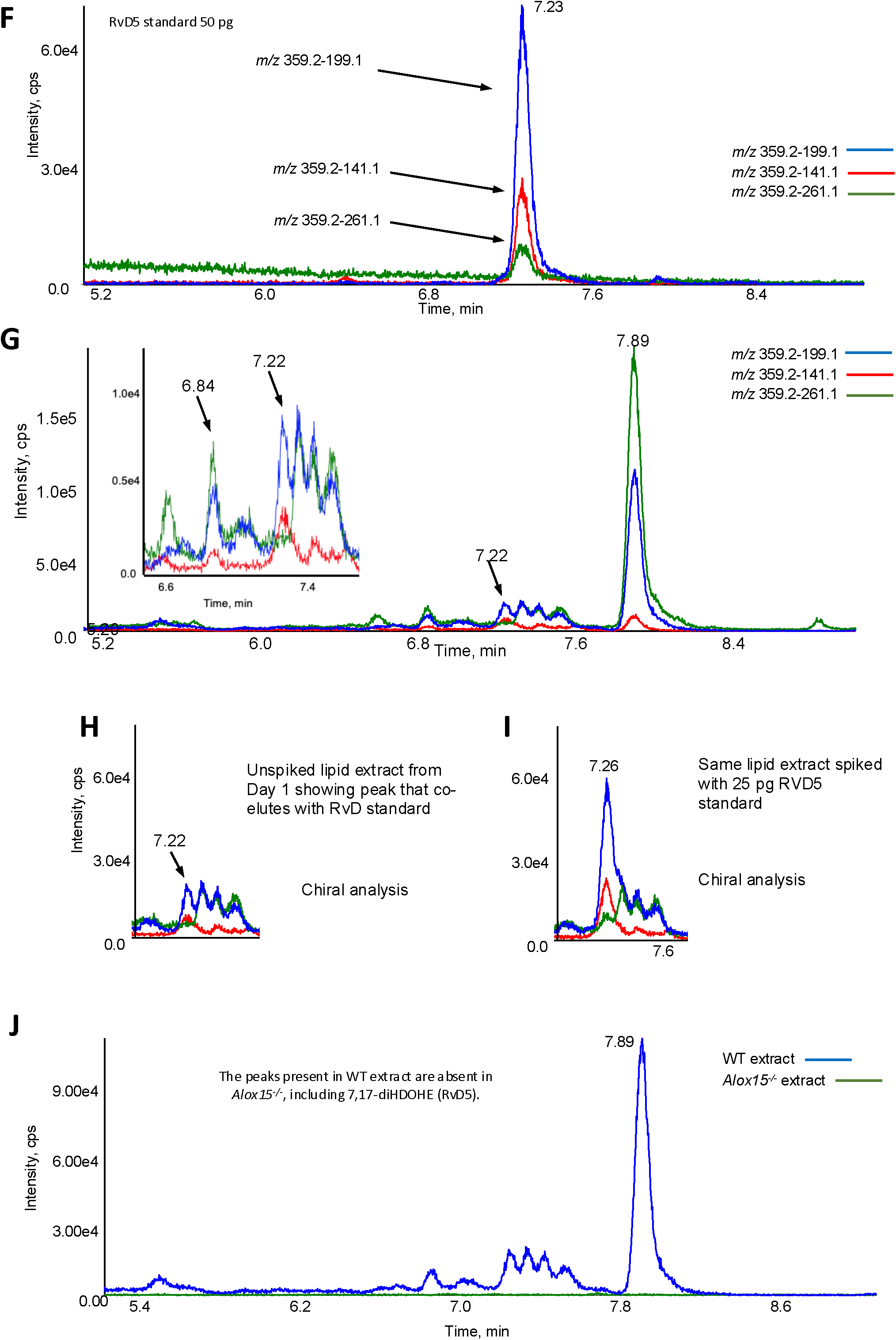
Reverse phase and chiral phase analysis of 7,17-diHDOHE suggests RvD5 along with additional isomers are present in WT mouse wounds at day 1. Lipid extracts from Day 1 wounds were pooled and analyzed using reverse and chiral phase LC/MS/MS as described in Methods. For reverse phase, the same method was used as for the oxylipin assay, but focusing on MRM transitions for RvD5, and removing scheduling. *Panels A-E. Reverse phase analysis of the synthetic RvD5 standard along with wound lipid extract from WT and Alox15^-/-^ mice.* Panels A,B,D show WT lipid extract, while Panel C shows the RvD5 standard. *Panels F-J. Chiral phase analysis of synthetic RvD5 standard along with wound lipid extract from WT and Alox15^-/-^ mice.* Panels G-J show wound extracts, while F shows the RvD5 standard.

**Supplementary Figure 13.**
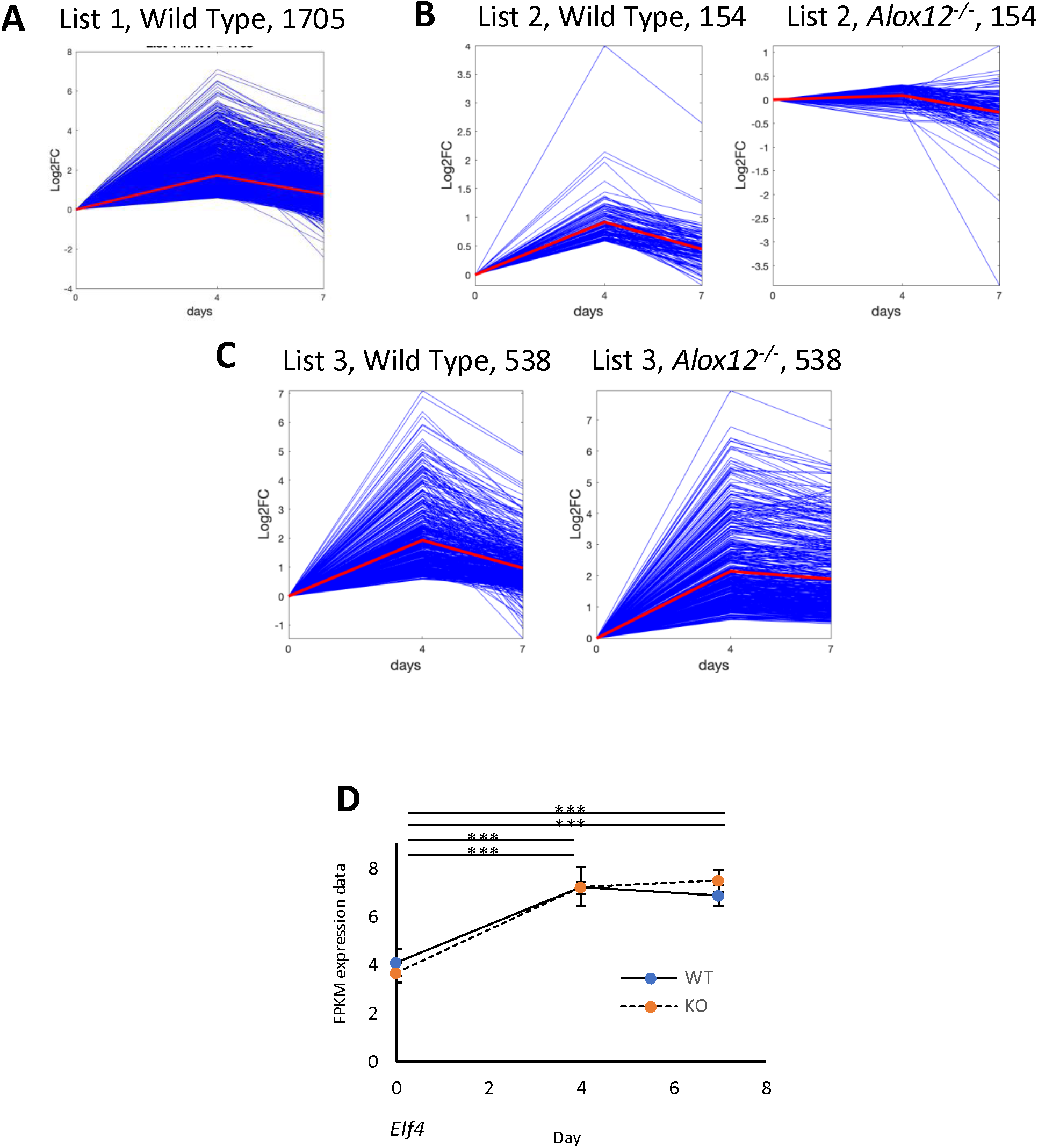
IPA temporal analysis shows altered regulation of several inflammatory pathways and induction of *Elf4*. *Panel A. List 1 represents 1705 transcripts in the WT condition* whose expression on Day 4 is significantly increased by > 50% compared to Day 0, i.e., with log2-fold change at least log2FC (1.5) and adjusted p-value < 0.05, but where log2FC values on Day 7 compared to Day 4 are reduced by > 25 %. *Panel B. List 2 represents a subset of List 1 consisting of 154 transcripts*, whose expression on Day 4 is not increased by > 25% compared to D0 in *Alox15^-/-^* wounds, i.e., with a maximum log2-fold change log2FC (1.25). Wilcoxon signed-rank test shows that the D4 data between the WT and *Alox15^-/-^* is significantly different. *Panel C.* List 3 comprises 538 transcripts from List 1 with a minimum log2-fold change log2FC (1.5) and adjusted p-value < 0.05 on D4 in *Alox15*^-/-^ wounds, whose log2FC values are reduced by < 25 % on Day 7 compared to Day 4 in *Alox15^-/-^* wounds. Wilcoxon signed-rank test shows that the D7 data between the WT and *Alox15^-/-^*conditions is significantly different. *Panel D. Elf4 is induced on wounding.* Gene expression data was normalized to its Day 0 mean value, and then expressed as fold-change (n = 3 – 4 per group). For all gene expression data, students t-test, followed by Benjamin Hochberg correction: * p <0.05, ** p < 0.01, *** p < 0.005.

**Supplementary Figure 14.**
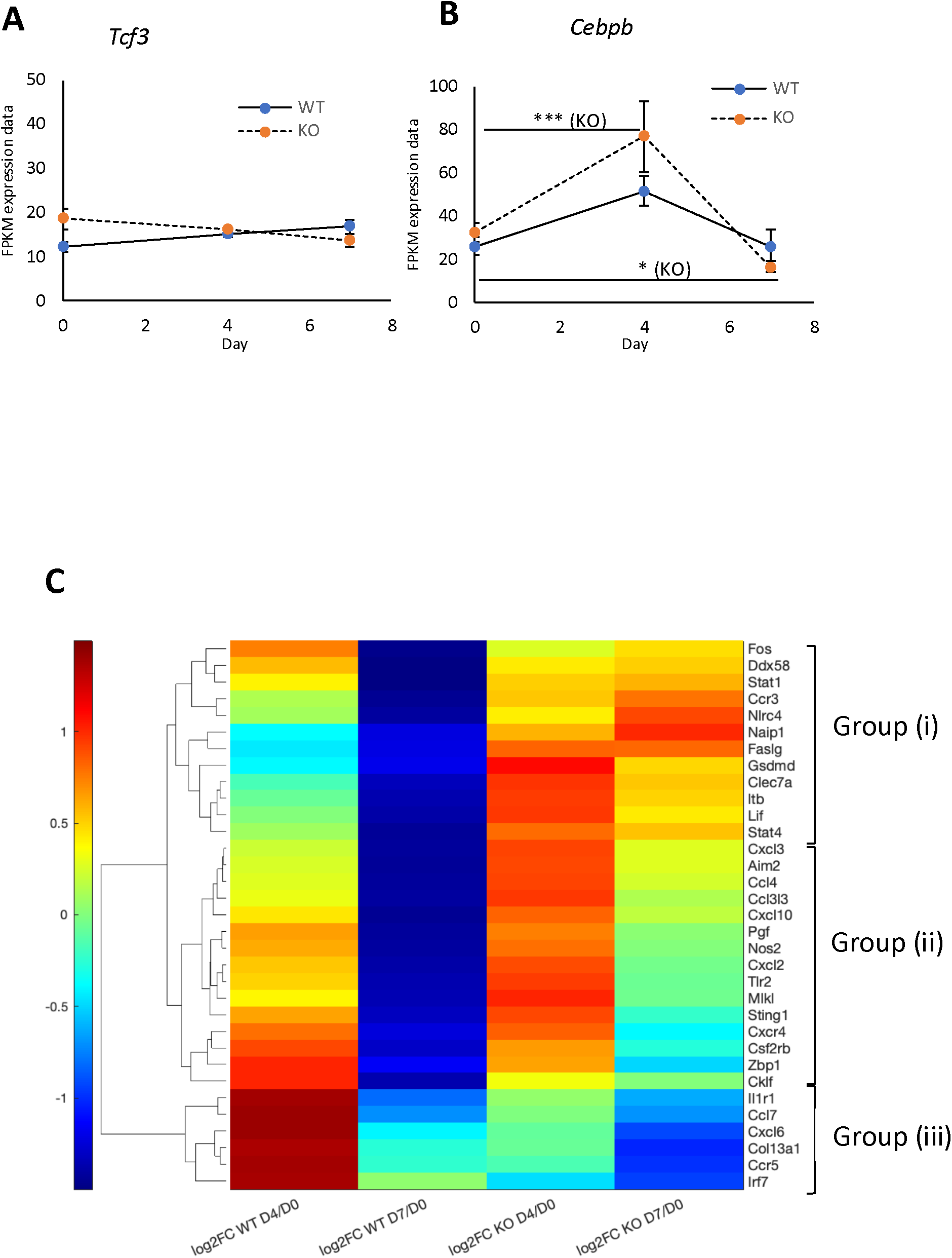
*Cebpb* is upregulated during wounding, but not Tcf3, while IPA identifies groups of genes with common behaviour that don’t resolve fully post wounding. *Panels A-C. Data from gene expression is shown for WT and Alox15^-/-^ wounds during the time course (n = 3 – 4 per group).* For all gene expression data, students t-test, followed by Benjamin Hochberg correction: * p <0.05, ** p < 0.01, *** p < 0.005. *Panel C. IPA analysis shows clusters of genes with similar behaviour, which don’t fully resolve.* Genes that do not fully resolve but behave in groups are plotted in this heatmap. Plotted are log2fold change data for wounds of the same strain comparing day 0 with either day 4 or day 7, within strain comparisons only.

